# New methods to calculate concordance factors for phylogenomic datasets

**DOI:** 10.1101/487801

**Authors:** Bui Quang Minh, Matthew W. Hahn, Robert Lanfear

## Abstract

We implement two measures for quantifying genealogical concordance in phylogenomic datasets: the gene concordance factor (gCF) and the novel site concordance factor (sCF). For every branch of a reference tree, gCF is defined as the percentage of “decisive” gene trees containing that branch. This measure is already in wide usage, but here we introduce a package that calculates it while accounting for variable taxon coverage among gene trees. sCF is a new measure defined as the percentage of decisive sites supporting a branch in the reference tree. gCF and sCF complement classical measures of branch support in phylogenetics by providing a full description of underlying disagreement among loci and sites. An easy to use implementation and tutorial is freely available in the IQ-TREE software package (http://www.iqtree.org).

---

Measures of branch support such as the bootstrap (Felsenstein 1985; Minh et al. 2013) and Bayesian posterior probability (Ronquist and Huelsenbeck 2003) are important for making robust inferences from phylogenetic trees. However, while these measures provide useful information about the statistical support for a given branch, neither captures the topological variation present in the underlying data (Kumar et al. 2012).

A complementary approach involves calculating the fraction of loci consistent with a particular branch, thus capturing underlying agreement and disagreement in the data. Various approaches have been suggested, including the gene concordance factor (gCF) (Baum 2007) — sometimes also referred to as the ‘gene support frequency’ (Gadagkar et al. 2005; Salichos and Rokas 2013) — and internode certainty (Salichos and Rokas 2013), with the former being the most widely used. The gCF describes for each branch in a reference tree the proportion of inferred single-locus trees that contain that branch. While intuitive, the gCF suffers from three limitations: first, there are few software implementations (Ané et al. 2007); second, while issues with incomplete taxon sampling of gene trees have been addressed by some authors (Smith et al. 2015; Kobert et al. 2016), these fixes are not available in implementations that allow for the calculation of gCF values; and third, low gCF values are hard to interpret because they may result from strongly supported discordance among individual gene trees or weak phylogenetic signal in individual loci.

Here, we resolve these issues by implementing the calculation of gCF while accounting for unequal taxon sampling in the popular IQ-TREE package (Nguyen et al. 2015; Minh et al. 2020), and by introducing the site concordance factor (sCF), a measure that estimates concordance at the level of individual sites. We argue, and show by detailed analyses of 10 empirical datasets, that concordance factors complement commonly used metrics like the bootstrap by providing additional information and insights about topological variation.

## Gene Concordance Factor (gCF)

For a given *unrooted* bifurcating reference tree, *T* (e.g. an estimate of the species tree), and a set of *unrooted* bifurcating input trees, *S* = {*T*_1_, …, *T*_*n*_} (e.g. individual gene trees), we can calculate the gCF for every internal branch *x* in *T*. Each gene tree, *T*_*i*_, must contain a subset of the taxa in *T*, but need not include all of the same taxa.

Each internal branch *x* of *T* is associated with four clades of *T* representing four taxon subsets, *A*, *B*, *C*, and *D*, such that the bipartition *A* ∪ *B*|*C* ∪ *D* corresponds to the branch *x* (Fig. 1). The reference tree *T* contains branch *x* by definition, but branch *x* may or may not be present in a gene tree of the same taxa (or a subset of them). Let *A*_*i*_, *B*_*i*_, *C*_*i*_, and *D*_*i*_ be the set of taxa in *A*, *B*, *C*, and *D* that are also present in *T*_*i*_. We say *T*_*i*_ is *decisive* for *x* if *A*_*i*_, *B*_*i*_, *C*_*i*_, *D*_*i*_ are non-empty (we use ‘decisive’ in the sense of (Sanderson et al. 2011), but note for clarity that Dell’Ampio et al. (2014) use this term in a different way): i.e. a gene tree is *decisive* as long as it contains at least one of the taxa in each of the taxon subsets *A*, *B*, *C*, and *D*. A *decisive* gene tree can potentially contain the branch *x*. We say *T*_*i*_ is *concordant* with *x* if *T*_*i*_ is *decisive* for *x* and the bipartition *A*_*i*_ ∪ *B*_*i*_|*C*_*i*_ ∪ *D*_*i*_ corresponds to a branch in *T*_*i*_. In other words, a gene tree is *concordant* with the reference tree if it could have contained branch *x* (i.e. it is *decisive*) and it does contain branch *x*. The gene concordance factor for branch *x* is now defined as:

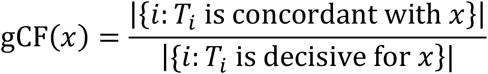

In other words, the gCF is the proportion of input trees decisive for *x* that are concordant with *x*; i.e. the proportion of all input trees that *could have* contained *x* that *do* contain *x*. We note that gCF is also used to calculate internode certainty and related measures (Salichos et al. 2014; Kobert et al. 2016).

**Figure 1.**
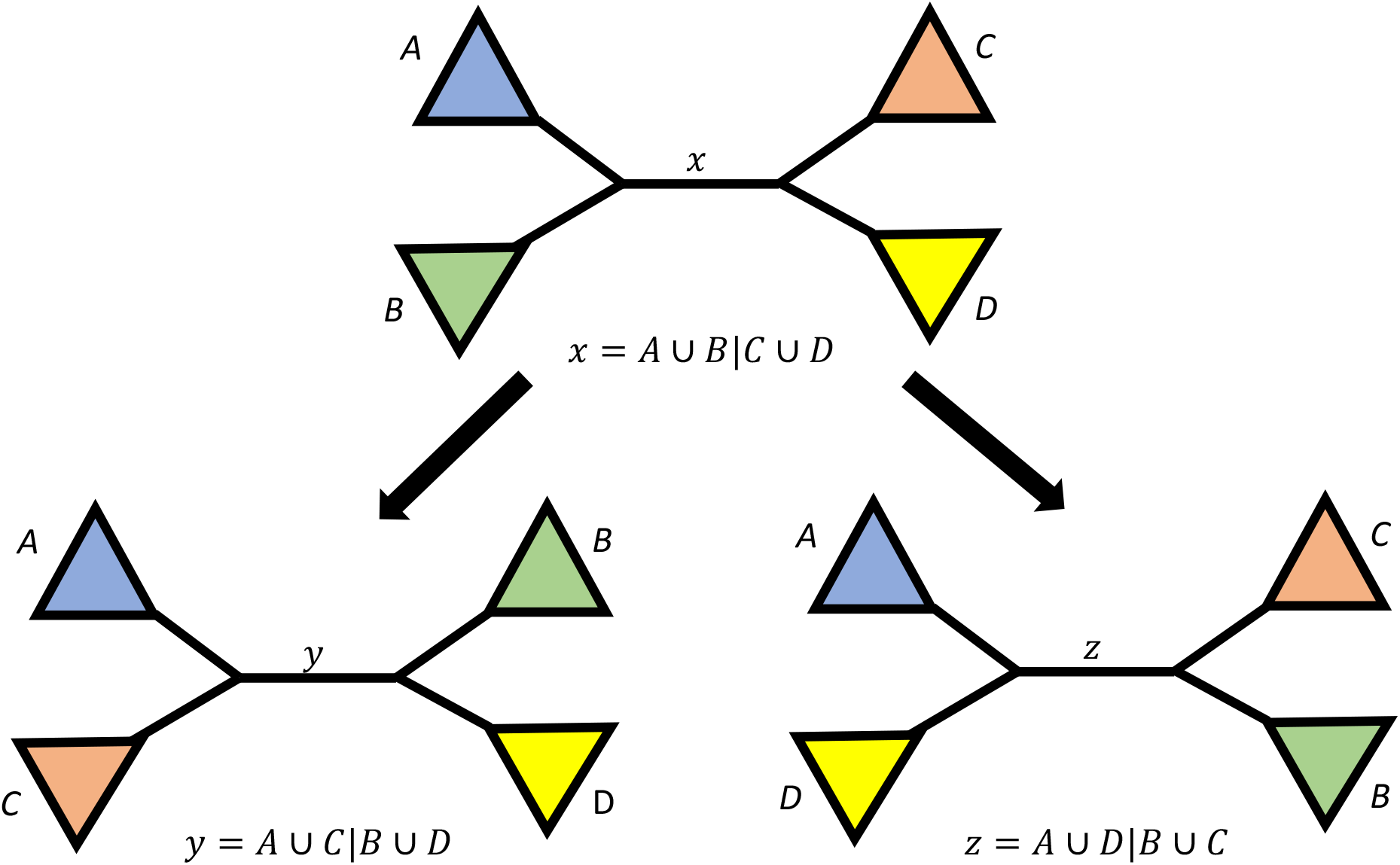
Schematic view of a bifurcating tree at an internal branch *x* with four surrounding sub-trees containing the taxon sets *A*, *B*, *C*, and *D*. Branch *x* corresponds to the bipartition *A* ∪ *B*|*C* ∪ *D*. In our definition, branch *x* is always present in the reference tree, but it may or may not be present in each of the input trees. By applying the two nearest neighbour interchanges (denoted as arrows) one can produce the two other trees that contain taxon sets *A*, *B*, *C*, and *D*, but with internal branches *y* and *z*.

To further understand *concordant* and *discordant* gene trees, we categorise the *discordant* input trees (i.e. those that do not contain branch *x*) into three groups, and calculate three *discordance* factors: gDF_1_, gDF_2_, and gDF_P_. The first two groups are obtained by applying a nearest neighbour interchange (NNI) around branch *x* to result in two alternative topologies (Fig. 1) with two branches not appearing in *T*: *y* = *A* ∪ *C*|*B* ∪ *D* and *z* = *A* ∪ *D*|*B* ∪ *C*. Accordingly, we then calculate discordance factors as gCF(*y*) = gDF_1_(*x*) and gCF(*z*) = gDF_2_(*x*). Some input trees may not contain any of the branches *x, y*, or *z*. This will occur for input trees in which one or more of clades *A*, *B*, *C*, or *D* are not monophyletic. We define the proportion of *decisive* input trees that fall into this category as gDF_P_(*x*), where the ‘P’ stands for ‘paraphyly’. The four proportions gCF(*x*), gDF_1_(*x*), gDF_2_(*x*), and gDF_P_(*x*) will sum to 1, as every *decisive* input tree must be included in one of the four categories.

## Site Concordance Factor (sCF)

To calculate the sCF for branch *x*, we randomly sample *m* quartets of taxa *q* = {*a*, *b*, *c*, *d*}, where *a*, *b*, *c*, and *d* are in *A*, *B*, *C*, and *D*, respectively. For each quartet *q*, we examine the sub-alignment of taxa *a*, *b*, *c*, *d*. For every site *j* in this alignment, we call *j decisive* for *x* if the characters *a*_*j*_, *b*_*j*_, *c*_*j*_, and *d*_*j*_ are all present and *j* is parsimony informative when restricted to this quartet of taxa. Decisive sites can be concordant or discordant with *x*. We say that site *j* is *concordant* with *x* if *a*_*j*_ = *b*_*j*_ ≠ *c*_*j*_ = *d*_*j*_ (i.e., *j* supports the bipartition {*a*, *b*}|{*c*, *d*}). The concordance factor for *q* is defined as:

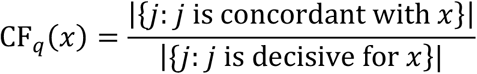

Since many such quartets exist around branch *x* (when sampling individual tips from within *A*, *B*, *C*, and *D*), we define sCF(*x*) as the mean CF_*q*_(*x*) over *m* random quartets:

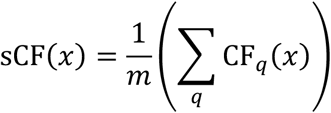

Thus, the sCF is the average proportion of sites decisive for *x* that are concordant with *x*. In effect, the sCF is a measure of concordance for sites that is directly comparable to the measure of concordance for single-locus trees provided by the gCF. Unlike related quartet approaches that are based on maximum likelihood tree inference and that calculate discordance at the level of the whole alignment (Strimmer and von Haeseler 1997; Pease et al. 2018), the sCF uses parsimony criteria to calculate discordance at the level of individual sites. We note that the sCF is closely related to the values derived from spectral analysis (Hendy and Penny 1993; Charleston 1998). Spectral analysis is a tree-independent method of investigating phylogenetic signal. In the simple case of binary data, such that each site corresponds to a bipartition of taxa, the spectral support for a single branch counts the number of sites that correspond to the bipartition defined by that branch. In the case of binary data, this is the same as the value CF_*q*_(*x*) that we define above. The sCF in this case differs from the spectral support insofar as it is calculated by averaging over a large set of CF_*q*_(*x*) values calculated by repeatedly subsampling quartets of taxa from an alignment.

Similarly to gene discordance factors, we also define the site discordance factors sCF(*y*) = sDF_1_(*x*) and sCF(*z*) = sDF_2_(*x*). There is no sDF_P_ category, because unlike the gDF values, every *decisive* site must contribute to one of the three proportions sCF(*x*), sDF_1_(*x*), or sDF_2_(*x*); any quartet of taxa resolves into exactly three tree topologies shown in Figure 1. In other words, the sum of sCF(*x*), sDF_1_(*x*), and sDF_2_(*x*) will always be 1.

## gCF and sCF for rooted trees

The above definitions of gCF and sCF apply only to unrooted trees. However, in the case that users have a rooted reference tree as well as rooted gene trees, we can extend the calculation of the gCF to allow us to calculate different gCF values on either side of the root. To do this we first add a virtual root node into the rooted reference tree *T*, resulting in an unrooted tree *T*’. Similarly, we convert the rooted gene trees *T*_*i*_ into unrooted trees *T*’_*i*_ by adding the same virtual root. This allows us to then follow the same procedure as above for calculating gCF values on unrooted trees, with the sole difference that the output is a rooted instead of an unrooted reference tree. In this case, it is possible to calculate gCF values on a rooted reference tree where the gCF values on either side of the root may differ. In all other cases (i.e. for sCF values, and if either the reference tree or the gene trees are unrooted), it is not possible to calculate different concordance factors on either side of the root.

## Implementation in IQ-TREE 2

We provide two new options, --gcf and --scf, in IQ-TREE version 2 (Minh et al. 2020) to compute gCF and sCF respectively. A tutorial for how to use these options is provided at http://www.iqtree.org/doc/Concordance-Factor. Both options can be combined in a single run, which will calculate the gCF and sCF for every branch in the input reference tree. While sCFs can be calculated on any alignment, the calculation of gCFs requires individual gene trees. We have therefore implemented a convenient option, -S, to specify a partition file or a directory of single-locus alignments in which IQ-TREE will infer separate trees for each partition or alignment (Minh et al. 2020).

IQ-TREE provides a suite of output files to assist users in understanding and investigating gCF and sCF values. It provides tree files that can be viewed in most tree viewers, which contain information on both the proportional data (gCF, gDF_1_, gDF_2_, gDF_P_; sCF, sDF_1_, sDF_2_) and their corresponding absolute count data (gCF_N, gDF_1__N, gDF_2__N, gDF_P__N; sCF_N, sDF_1__N, sDF_2__N), as well as the number of decisive genes and sites for each branch (gN and sN, respectively). It also provides a tree file that combines information on the gCF, sCF, and any bootstrap values that have been calculated (e.g. Figure 2). In addition to this, IQ-TREE provides a ‘.cf.stat’ file that contains all 16 concordance and discordance values listed above for every branch in the reference tree in a machine-readable tabulated format, and through the --cf-verbose option it provides tabulated files that detail for every branch in the reference tree whether each gene tree was concordant with that branch (the ‘.cf.stat_tree’ file produced from a --gcf analysis), and the average number of sites in each locus that were concordant with that branch (the ‘.cf.stat_loci’ file produced from a --scf analysis). Together these output files provide both a convenient overview of the data, and the opportunity to understand in much more detail the extent to which each locus is concordant or discordant with the reference tree.

**Figure 2.**
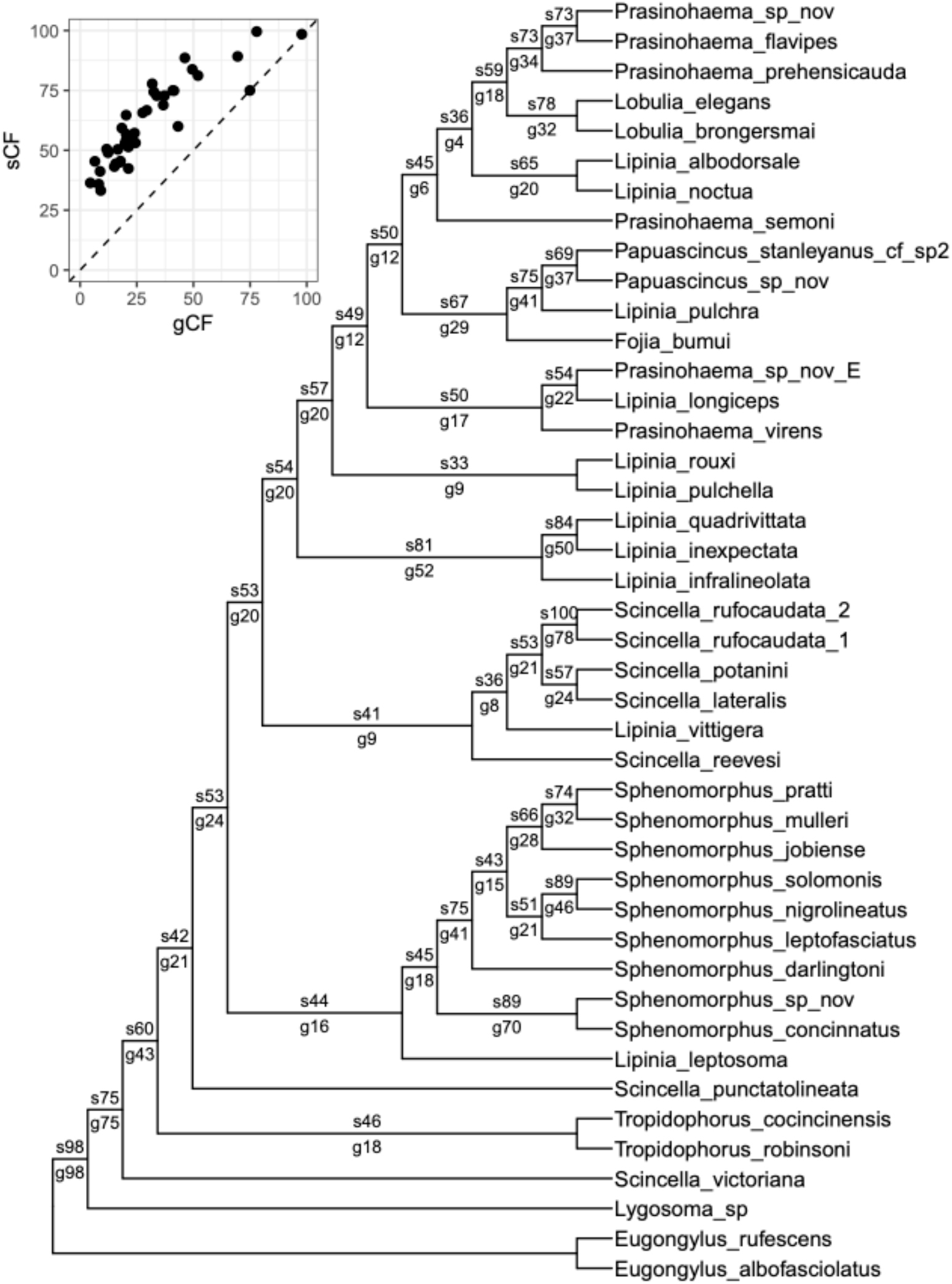
An example of concordance factors on a dataset of lizards (Rodriguez et al. 2018). A cladogram is shown to facilitate the plotting of concordance factors on branches. Numbers on each branch show the site concordance factor (sCF) above the branch (e.g. s73) and the gene concordance factor (gCF) below the branch (e.g. g37). Bootstrap values are 100% on every branch, except for the branch leading to *Lipinia rouxi* and *L. pulchella*, which has a bootstrap value of 62%. The inset shows a scatter plot of gCF values against sCF values for all branches, revealing the large range of gCF and sCF values as well as the fact that for this dataset sCF values are always at least as large as gCF values. This is likely because of the short length of the UCE loci used to infer gene trees (Rodriguez et al. 2018).

## Application to empirical datasets

To demonstrate the use of gCF and sCF values, we first analysed a dataset containing 3,220 ultra-conserved elements (UCEs) from lizards, totalling 1,301,107 bp for 43 species (Rodriguez et al. 2018). To do this, we estimated a concatenated maximum likelihood (ML) tree, 3,220 UCE trees, and the gCF, sCF, and bootstrap values with IQ-TREE (Figure 2).

To investigate the relationships between dataset size, concordance factors, and bootstrap values, we also analysed a collection of nine additional empirical phylogenomic datasets that represent a range of clades and data types (Table 1). For each dataset, we performed 20 analyses across a 20-fold range of dataset sizes: the first analysis included 10 randomly selected loci from the complete dataset, and each subsequent analysis added 10 more randomly selected loci up to a maximum of 200 loci. For each such analysis, we calculated the gCF, sCF, and bootstrap values for every branch of the concatenated ML tree estimated from the 200-locus dataset, with the same approach as above. For the lizard dataset, we calculated both standard bootstrap (StdBoot) values (Felsenstein 1985) and ultrafast bootstrap (UFBoot) values (Hoang et al. 2018). As expected (Minh et al. 2013), we observed that StdBoot and UFBoot values were very similar (Figure 3B, supplementary figure 8D); therefore, because of the high computational cost of calculating StdBoot values, we calculated only UFBoot values for the remaining nine datasets. Since the results of all 10 analyses are very similar, we focus here on the lizard dataset, and present the results for the other 9 datasets in the supplementary figures.

**Table 1.** The five DNA and five amino acid (AA) datasets analysed in this study.

| No | Reference | Type | Clade | Taxa | Loci | Sites |
| --- | --- | --- | --- | --- | --- | --- |
| 1 | (Ballesteros and Sharma 2019) | AA | Chelicerata | 53 | 3,534 | 1,484,206 |
| 2 | (Branstetter et al. 2017) | DNA | Aculeata | 187 | 807 | 183,747 |
| 3 | (Cannon et al. 2016) | DNA | Metazoa | 78 | 424 | 89,792 |
| 4 | (Jarvis et al. 2015) | AA | Aves | 52 | 8,295 | 4,519,041 |
| 5 | (Misof et al. 2014) | AA | Insecta | 144 | 2,868 | 595,033 |
| 6 | (Ran et al. 2018) | AA | Spermatophyta | 38 | 1,308 | 432,014 |
| 7 | (Ran et al. 2018) | DNA | Spermatophyta | 38 | 3,924 | 1,296,042 |
| 8 | (Rodriguez et al. 2018) | DNA | Prasinohaema | 43 | 3,220 | 1,301,107 |
| 9 | (Wu et al. 2018) | AA | Mammalia | 90 | 5,162 | 3,050,199 |
| 10 | (Wu et al. 2018) | DNA | Mammalia | 90 | 15,486 | 9,150,597 |

**Figure 3.**
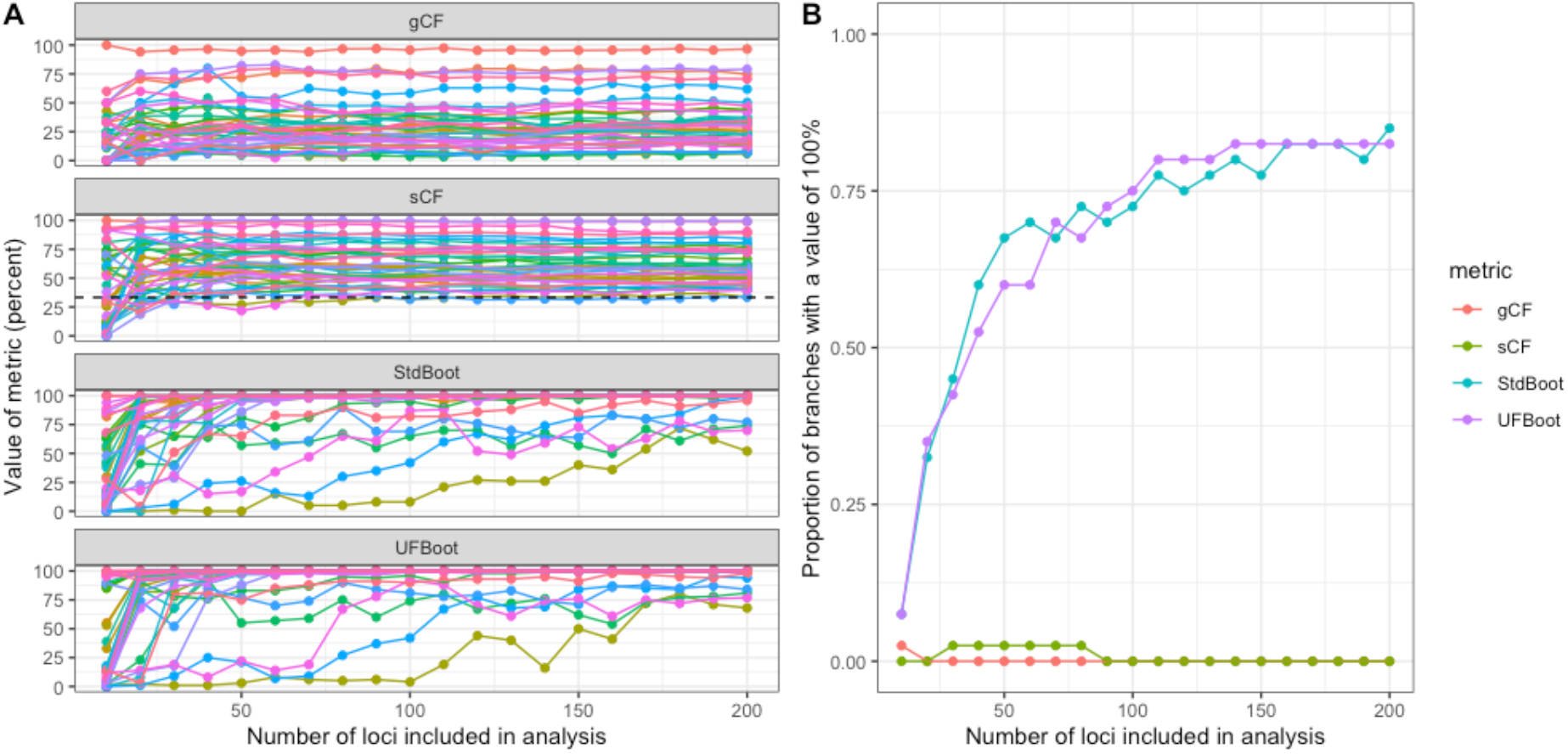
Concordance factors remain relatively stable as loci are added to an analysis, while bootstrap values continue to increase towards 100%. A) The results of 20 reanalyses of the lizard dataset, each of which adds a further ten loci to the analysis, up to a maximum of 200 loci (x-axis). Each coloured line represents a different branch in the tree. The dashed line on the sCF panel shows a value of 33%. B) For each of the four metrics considered here (represented by different coloured lines), the number of loci included in the analysis affects the proportion of the branches that have a value of 100%.

Reproducible analyses are provided in the supplementary material available at https://doi.org/10.5281/zenodo.1949287.

## Interpretation of gCF and sCF values

Our analyses highlight a number of important differences between concordance factors and bootstrap values. First, concordance factors of any type tend to be much lower than bootstrap values or other measures of statistical support for branches in the species tree (e.g. Fig 3, supplementary figures 1 to 10). This is because bootstrap values measure the sampling variance in support of a focal branch, while gCF and sCF values measure the underlying variance in support of that branch at the gene- and site-level respectively. In other words, resampled datasets may always return the same tree (i.e. 100% bootstrap support), even though incomplete lineage sorting or other processes that lead to genealogical discordance are at work (e.g. gCF and sCF values ≪100%). Of particular note is that a very high bootstrap value (e.g. of 100%) does not predict or require a similarly high concordance factor. For example, although all but one bootstrap value on the tree in Figure 1 is 100%, the smallest gCF and sCF values are 4.5% and 33.2% respectively. This pattern is repeated across all 200 analyses we performed on all 10 empirical datasets (Figure 3, supplementary figures 1 to 10).

Because they measure different things, concordance factors and bootstrap values are affected very differently by the addition of loci to a dataset (Figure 3, supplementary figures 1 to 10). In all of the 10 empirical datasets that we analysed, adding loci to the dataset tended to increase the bootstrap values of individual branches (e.g. Figure 3A, see also supplementary figures) and concomitantly the proportion of branches with a bootstrap value of 100% (Figure 3B, see also supplementary information). This is expected: the bootstrap is effectively measuring the standard error of the mean in a dataset (where the mean represents the tree inferred from the full dataset), and this standard error will go down with more samples. The same is not true of concordance factors, which display some estimation error when datasets are small (e.g. 30 or fewer loci in Figure 3), but subsequently remain almost completely insensitive to the addition of loci to the dataset (Figure 3, supplementary figures 1-10). This is also expected: any measure of the underlying variance (or standard deviation) in a distribution will have some estimation error when the sample size is small, but should not change monotonically as the sample size increases. These differences highlight the complementary information that can be gained from calculating both bootstrap values and concordance factors for individual branches. One measure is not better or worse than the other, rather, concordance factors provide useful information that bootstraps do not, and vice versa. Note that bootstrap values or posterior probabilities calculated by resampling gene trees (e.g., Sayyari and Mirarab 2016) have the same behaviour as the site-wise bootstrap carried out here, and are also not equivalent to concordance factors.

In principle, both gCF and sCF values can range from 0% (no genes / sites are concordant with the focal branch) to 100% (all genes / sites are concordant with the focal branch). In practice however, as exemplified in Figure 3 and supplementary figures 1 to 10, empirical gCF values tend to range from 0% to 100%, while empirical sCF values are rarely lower than 33% (represented by the dashed line in the sCF panel of Figure 3A). This is due to an important underlying difference in the way that the two values are calculated. The sCF is calculated from quartets, so a single site can only support one of three topologies (Figure 1). Because of this, if there is no consistent information in an alignment (e.g. if a long alignment were generated at random) we expect a roughly equal proportion of sites supporting each of the three trees, leading to an sCF value of approximately 33% (for the same reason, sCF values for very long branches will approach 33% due to saturation). The same is not true for gCF values, because a gene tree can support not only the three possible relationships shown in Figure 1, but any other relationship in which one or more of clades *A*, *B*, *C*, or *D* is not monophyletic. The higher the number of gene trees in this latter group, the closer the gCF value will be to 0%. Because of this, we should expect gCF values to be particularly low when gene trees are estimated from alignments with limited information or where branch *x* is extremely short; in such cases either technical or biological processes may increase the proportion of gene trees that fail to recover the monophyly of clades *A*, *B*, *C*, or *D* found in the reference tree. Missing data may also impact gCF and sCF values, however, the relationship here is less clear. The requirement that a gene tree or a site is decisive (i.e. could in principle contain branch *x*, see above) should limit the impact of missing data on gCF and sCF estimates. Nevertheless, since phylogenetic estimates are known to worsen as the proportion of missing data increases (e.g., Roure et al. 2013; Xi et al. 2016), it is plausible that concordance factors may systematically decrease as the proportion of missing data increases.

Finally, cases where the sCF value is lower than 33% may be of particular interest. These cases are, by definition, those in which maximum parsimony (MP) would favour a different resolution of a split found in the reference tree. If the reference tree was calculated from any method other than MP, there are at least two explanations for an sCF value lower than 33%. First, the branch of interest may be in an area of parameter space in which high levels of incomplete lineage sorting are known to mislead concatenated ML analyses (Kubatko and Degnan 2007) but not MP analyses (Mendes and Hahn 2018). Misleading reference trees may therefore be produced by either concatenated ML of the entire dataset, or by gene tree methods that use shorter sets of concatenated loci as their input (because most protein-coding genes are themselves made up of multiple topologies). Second, and more generally, there are multiple reasons why likelihood, Bayesian, or gene tree methods for producing a species tree will differ from MP resolutions. For instance, the branch of interest may be unduly affected by a small number of highly influential sites in a concatenated ML analysis (Shen et al. 2017). In this case, the influential sites can have an outsized influence on the ML resolution of a split because they have extreme differences in likelihood between different resolutions of that split. Because MP does not account for likelihood differences—it instead weights all sites equally—MP analyses remain unaffected by such outliers. Thus, cases in which the sCF is much lower than 33% may merit further investigation.

We hope that the user-friendly implementation of gene- and site-concordance factors in IQ-TREE will assist researchers in gaining additional insights into their phylogenetic reconstructions. In particular, we encourage phylogeneticists to calculate both bootstrap values and concordance factors for the branches on their trees, as the two measures provide complementary information that may help to improve the accuracy of our interpretations of phylogenetic reconstructions. Indeed, the use of concordance factors may help to alleviate the commonly cited problem in phylogenomics that bootstrap values provide relatively little information when they are all 100% (Kumar et al. 2012).

## Acknowledgements

We thank Zachary Rodriguez for providing the lizard dataset, Cécile Ané for suggesting the analogy with standard error and standard deviation, and the associate editor and two reviewers for helpful suggestions. This work was supported by National Science Foundation grant DEB-1936187 to MWH, an Australian National University Futures grant to RML, and an Australian Research Council grant DP200103151 to RML, BQM, and MWH.

## Supplementary Materials

Data are available at https://doi.org/10.5281/zenodo.1949287.

Supplementary figures are available from the journal website.

## References

Ané C, Larget B, Baum DA, Smith SD, Rokas A. 2007. Bayesian estimation of concordance among gene trees. Mol Biol Evol 24:412–426.

Ballesteros JA, Sharma PP. 2019. A Critical Appraisal of the Placement of Xiphosura (Chelicerata) with Account of Known Sources of Phylogenetic Error. Syst Biol 68:896–917.

Baum DA. 2007. Concordance trees, concordance factors, and the exploration of reticulate genealogy. Taxon 56:417–426.

Branstetter MG, Danforth BN, Pitts JP, Faircloth BC, Ward PS, Buffington ML, Gates MW, Kula RR, Brady SG. 2017. Phylogenomic Insights into the Evolution of Stinging Wasps and the Origins of Ants and Bees. Curr Biol 27:1019–1025.

Cannon JT, Vellutini BC, Smith J, 3rd, Ronquist F, Jondelius U, Hejnol A. 2016. Xenacoelomorpha is the sister group to Nephrozoa. Nature 530:89–93.

Charleston MA. 1998. Spectrum: spectral analysis of phylogenetic data. Bioinformatics 14:98–99.

Dell’Ampio E, Meusemann K, Szucsich NU, Peters RS, Meyer B, Borner J, Petersen M, Aberer AJ, Stamatakis A, Walzl MG, et al. 2014. Decisive data sets in phylogenomics: Lessons from studies on the phylogenetic relationships of primarily wingless insects. Mol Biol Evol 31:239–249.

Felsenstein J. 1985. Confidence limits on phylogenies: an approach using the bootstrap. Evolution 39:783–791.

Gadagkar SR, Rosenberg MS, Kumar S. 2005. Inferring species phylogenies from multiple genes: Concatenated sequence tree versus consensus gene tree. J Exp Zool Part B 304b:64–74.

Hendy MD, Penny D. 1993. Spectral analysis of phylogenetic data. Journal of Classification 10:5–24.

Hoang DT, Chernomor O, von Haeseler A, Minh BQ, Le SV. 2018. UFBoot2: Improving the ultrafast bootstrap approximation. Mol Biol Evol 35:518–522.

Jarvis ED, Mirarab S, Aberer AJ, Li B, Houde P, Li C, Ho SY, Faircloth BC, Nabholz B, Howard JT, et al. 2015. Phylogenomic analyses data of the avian phylogenomics project. Gigascience 4:4.

Kobert K, Salichos L, Rokas A, Stamatakis A. 2016. Computing the internode certainty and related measures from partial gene trees. Mol Biol Evol 33:1606–1617.

Kubatko LS, Degnan JH. 2007. Inconsistency of phylogenetic estimates from concatenated data under coalescence. Syst Biol 56:17–24.

Kumar S, Filipski AJ, Battistuzzi FU, Pond SLK, Tamura K. 2012. Statistics and truth in phylogenomics. Mol Biol Evol 29:457–472.

Mendes FK, Hahn MW. 2018. Why concatenation fails near the anomaly zone. Syst Biol 67:158–169.

Minh BQ, Nguyen MAT, von Haeseler A. 2013. Ultrafast approximation for phylogenetic bootstrap. Mol Biol Evol 30:1188–1195.

Minh BQ, Schmidt HA, Chernomor O, Schrempf D, Woodhams MD, von Haeseler A, Lanfear R. 2020. IQ-TREE 2: New models and efficient methods for phylogenetic inference in the genomic era. Mol Biol Evol in press.

Misof B, Liu S, Meusemann K, Peters RS, Donath A, Mayer C, Frandsen PB, Ware J, Flouri T, Beutel RG, et al. 2014. Phylogenomics resolves the timing and pattern of insect evolution. Science 346:763–767.

Nguyen LT, Schmidt HA, von Haeseler A, Minh BQ. 2015. IQ-TREE: A fast and effective stochastic algorithm for estimating maximum-likelihood phylogenies. Mol Biol Evol 32:268–274.

Pease JB, Brown JW, Walker JF, Hinchliff CE, Smith SA. 2018. Quartet Sampling distinguishes lack of support from conflicting support in the green plant tree of life. Am J Bot 105:385–403.

Ran JH, Shen TT, Wang MM, Wang XQ. 2018. Phylogenomics resolves the deep phylogeny of seed plants and indicates partial convergent or homoplastic evolution between Gnetales and angiosperms. P Roy Soc B-Biol Sci 285.

Rodriguez ZB, Perkins SL, Austin CC. 2018. Multiple origins of green blood in New Guinea lizards. Sci Adv 4:eaao5017.

Ronquist F, Huelsenbeck JP. 2003. MrBayes 3: Bayesian phylogenetic inference under mixed models. Bioinformatics 19:1572–1574.

Roure B, Baurain D, Philippe H. 2013. Impact of missing data on phylogenies inferred from empirical phylogenomic data sets. Mol Biol Evol 30:197–214.

Salichos L, Rokas A. 2013. Inferring ancient divergences requires genes with strong phylogenetic signals. Nature 497:327–331.

Salichos L, Stamatakis A, Rokas A. 2014. Novel Information Theory-Based Measures for Quantifying Incongruence among Phylogenetic Trees. Mol Biol Evol 31:1261–1271.

Sanderson MJ, McMahon MM, Steel M. 2011. Terraces in phylogenetic tree space. Science 333:448–450.

Sayyari E, Mirarab S. 2016. Fast Coalescent-Based Computation of Local Branch Support from Quartet Frequencies. Mol Biol Evol 33:1654–1668.

Shen XX, Hittinger CT, Rokas A. 2017. Contentious relationships in phylogenomic studies can be driven by a handful of genes. Nat Ecol Evol 1:126.

Smith SA, Moore MJ, Brown JW, Yang Y. 2015. Analysis of phylogenomic datasets reveals conflict, concordance, and gene duplications with examples from animals and plants. BMC Evol Biol 15:150.

Strimmer K, von Haeseler A. 1997. Likelihood-mapping: a simple method to visualize phylogenetic content of a sequence alignment. Proc Natl Acad Sci U S A 94:6815–6819.

Wu S, Edwards S, Liu L. 2018. Genome-scale DNA sequence data and the evolutionary history of placental mammals. Data Brief 18:1972–1975.

Xi Z, Liu L, Davis CC. 2016. The Impact of Missing Data on Species Tree Estimation. Mol Biol Evol 33:838–860.

