## Supplementary Figures for "New methods to calculate concordance factors for phylogenomic datasets"

### Supplementary Figure 1A

Dataset: Ballesteros_2019
A scatterplot of gCF vs sCF values, coloured by UFBoot values. Each point represents a single branch in the tree calculated from the 200 locus dataset. Each panel represents an analysis in which the concordance factors and bootstraps were calculated from the number of loci specified in the panel header. The dashed line is the 1:1 line

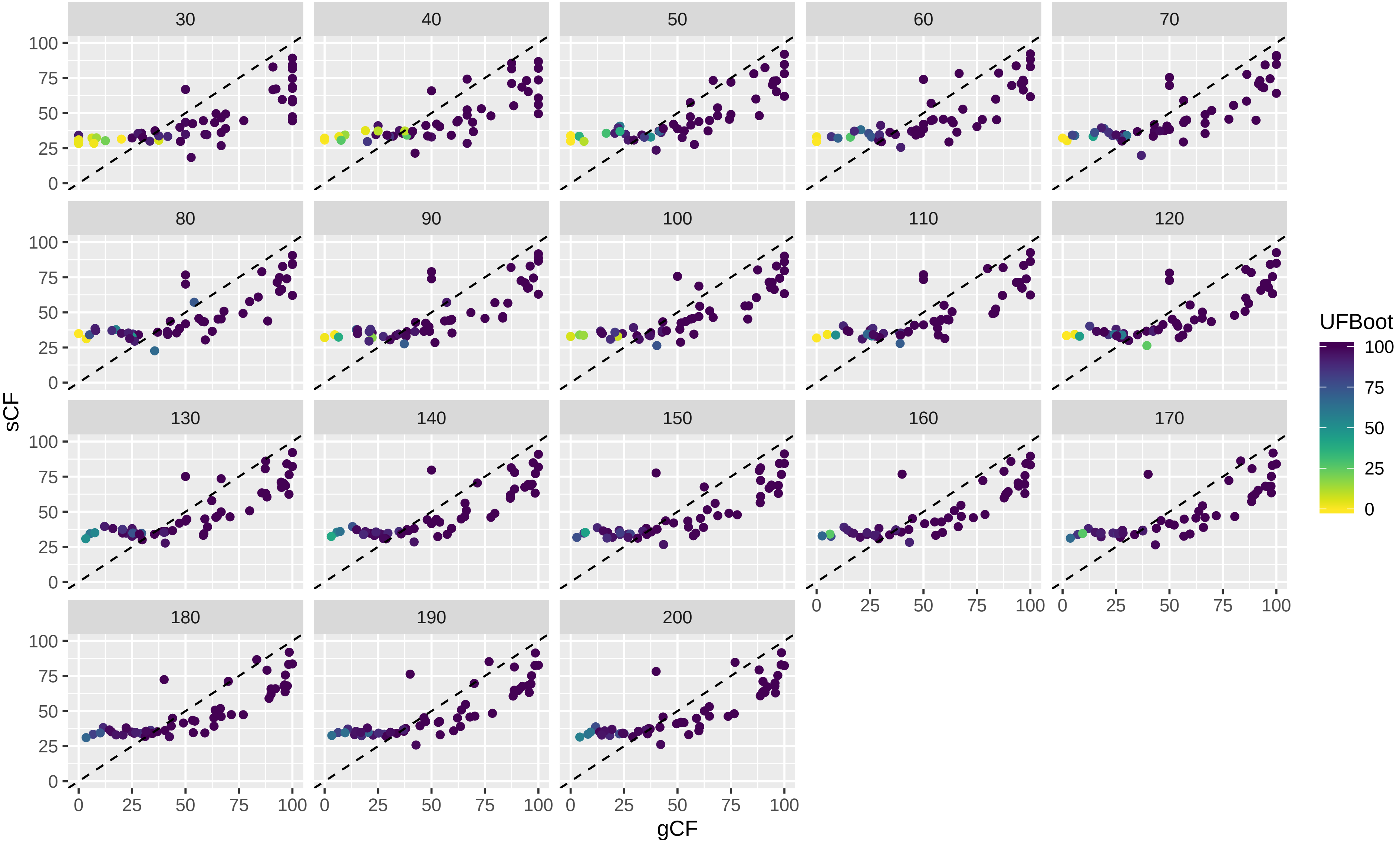

Supplementary Figure 1B

Dataset: Ballesteros_2019
The value of each variable (where the variable name is shown in the panel header) by number of loci used in the analysis.
Each coloured line represents a single branch in the tree estimated from the 200 locus dataset.

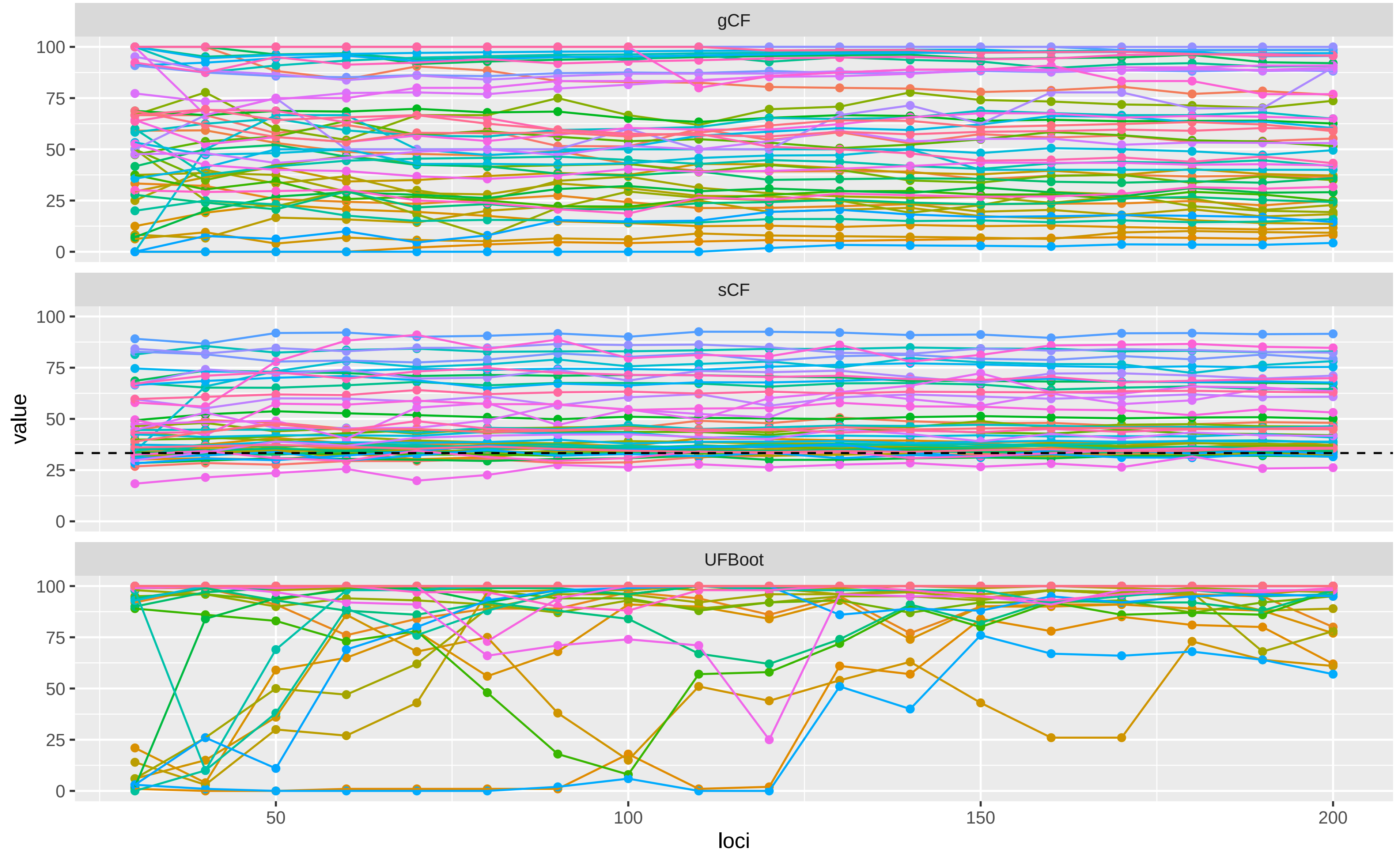

Supplementary Figure 1C

Dataset: Ballesteros_2019

The proportion of CF or Bootstrap values that are 100% (y axis) versus the number of loci used in the analysis (x axis).

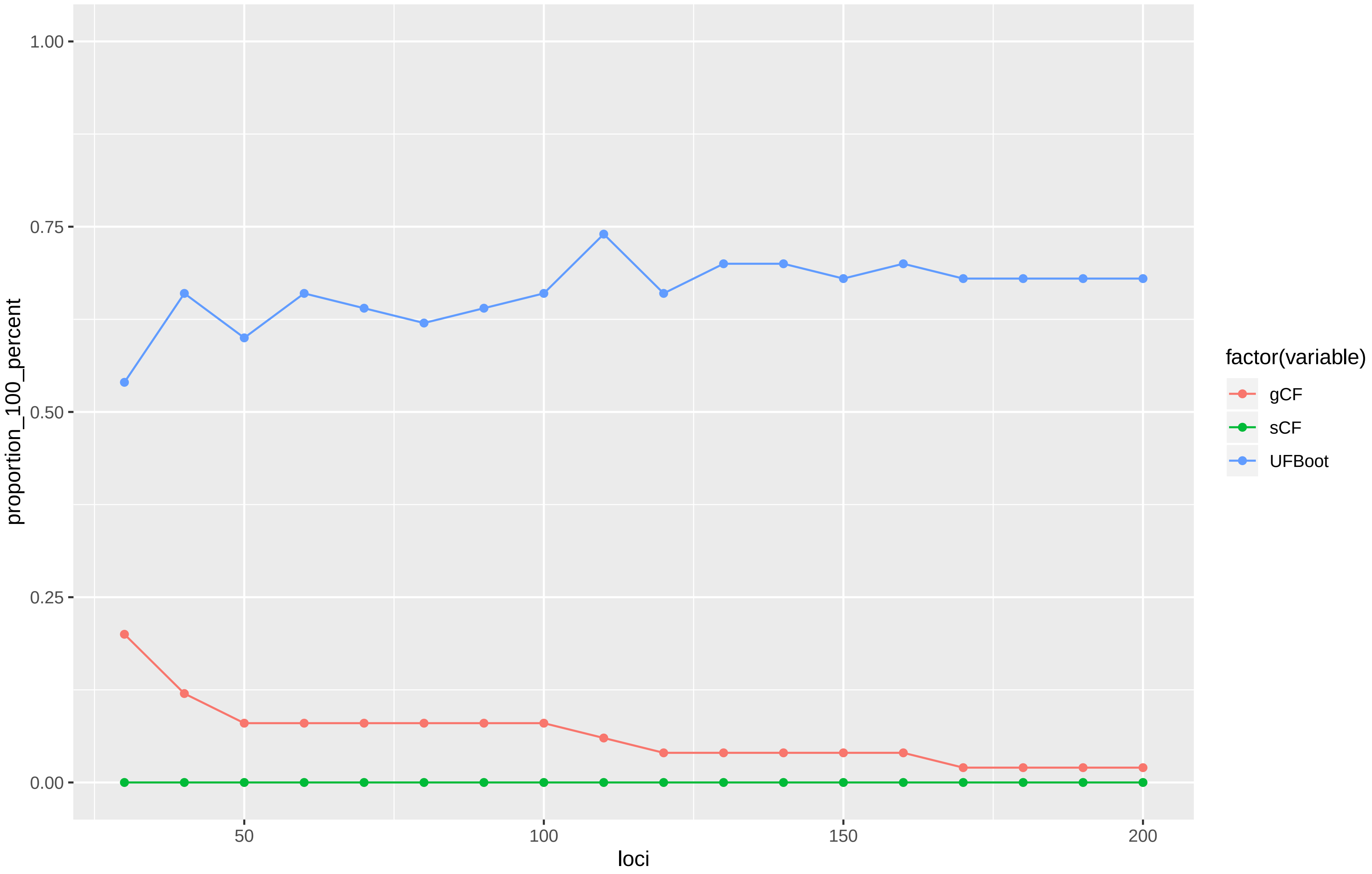

Supplementary Figure 2A

Dataset: Branstetter_2017
A scatterplot of gCF vs sCF values, coloured by UFBoot values. Each point represents a single branch in the tree calculated from the 200 locus dataset. Each panel represents an analysis in which the concordance factors and bootstraps were calculated from the number of loci specified in the panel header. The dashed line is the 1:1 line

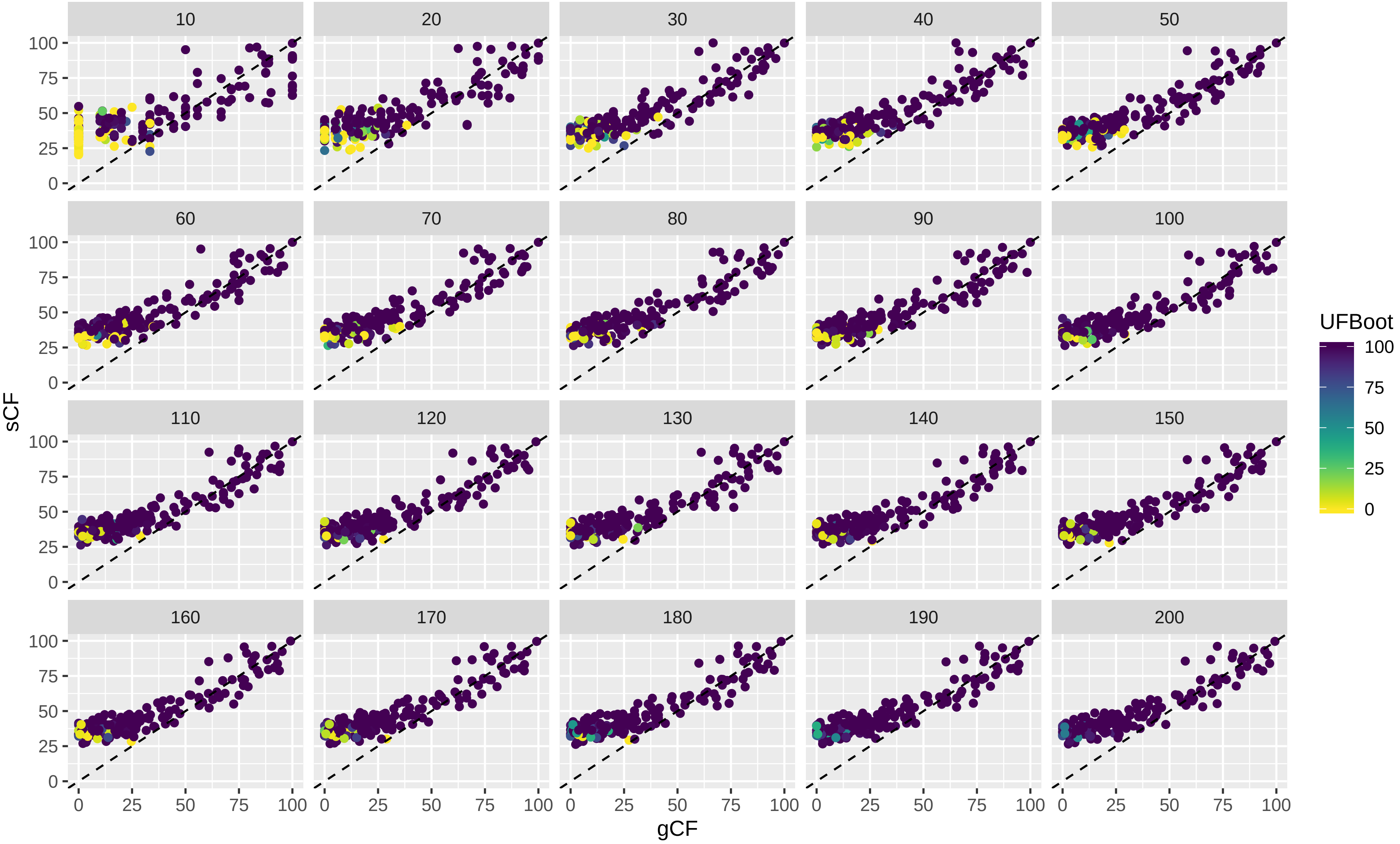

Supplementary Figure 2B

Dataset: Branstetter_2017
The value of each variable (where the variable name is shown in the panel header) by number of loci used in the analysis.
Each coloured line represents a single branch in the tree estimated from the 200 locus dataset.

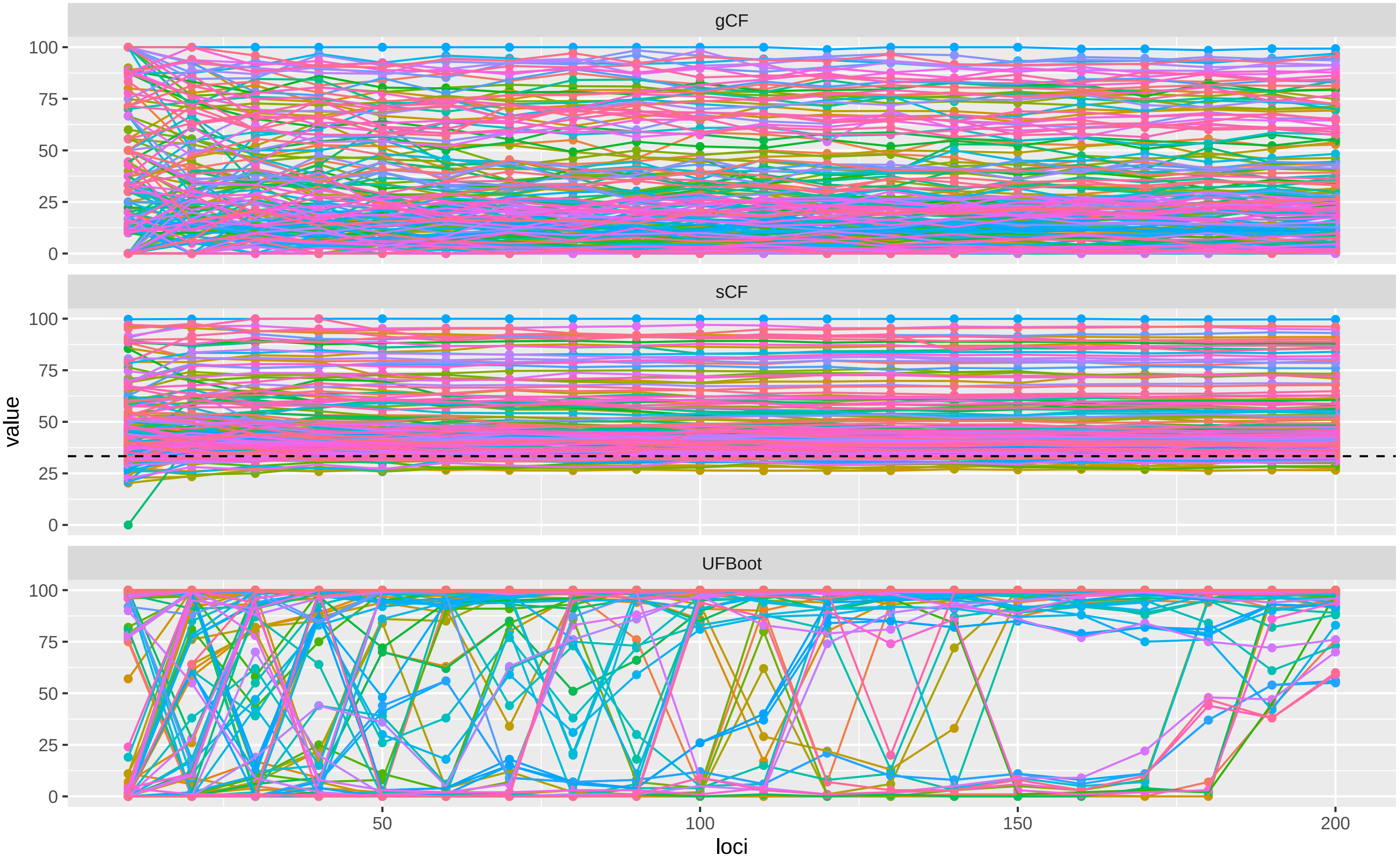

Supplementary Figure 2C

Dataset: Branstetter_2017

The proportion of CF or Bootstrap values that are 100% (y axis) versus the number of loci used in the analysis (x axis).

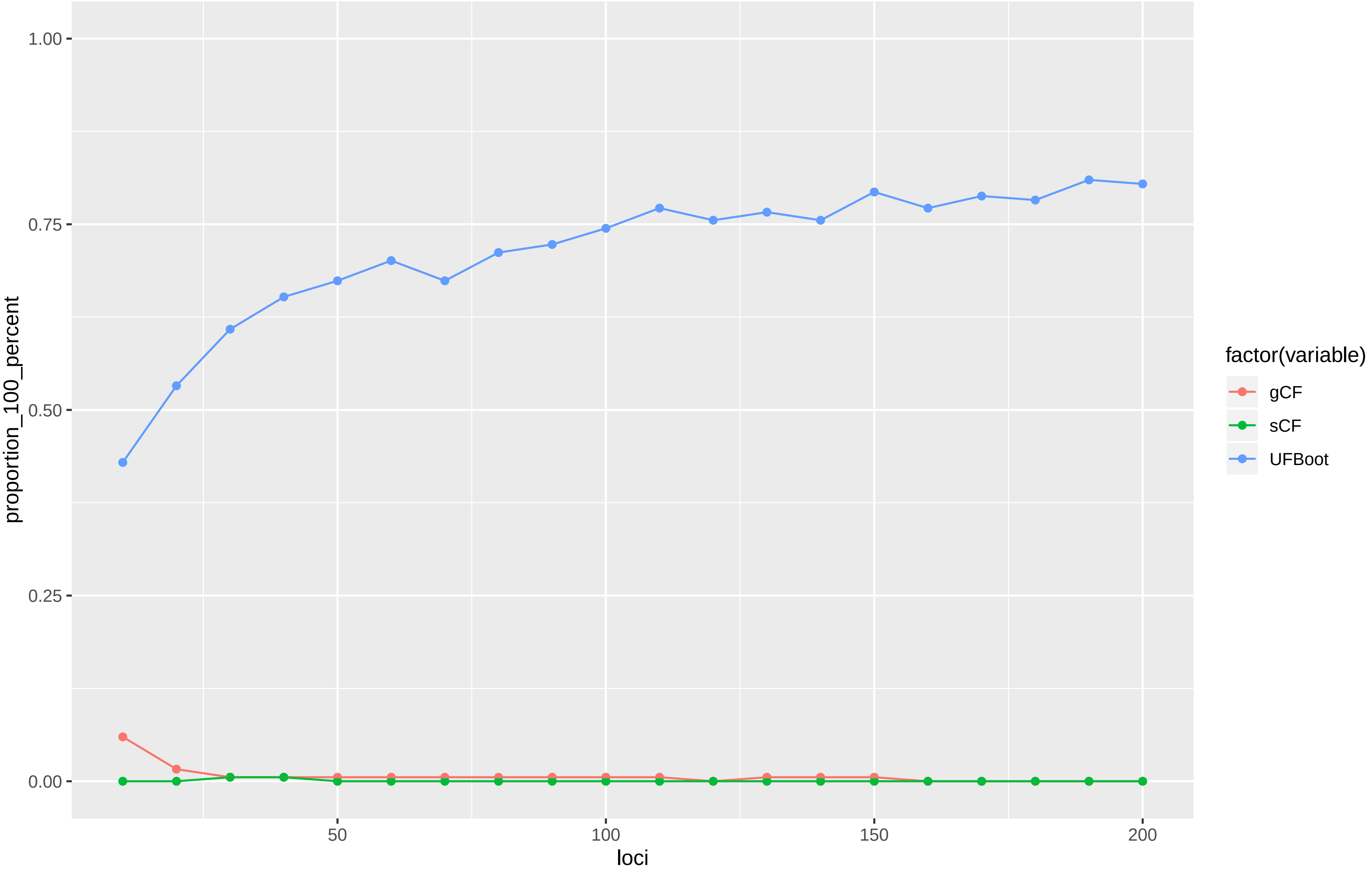

Supplementary Figure 3A

Dataset: Cannon_2016

A scatterplot of gCF vs sCF values, coloured by UFBoot values. Each point represents a single branch in the tree calculated from the 200 locus dataset. Each panel represents an analysis in which the concordance factors and bootstraps were calculated from the number of loci specified in the panel header. The dashed line is the 1:1 line

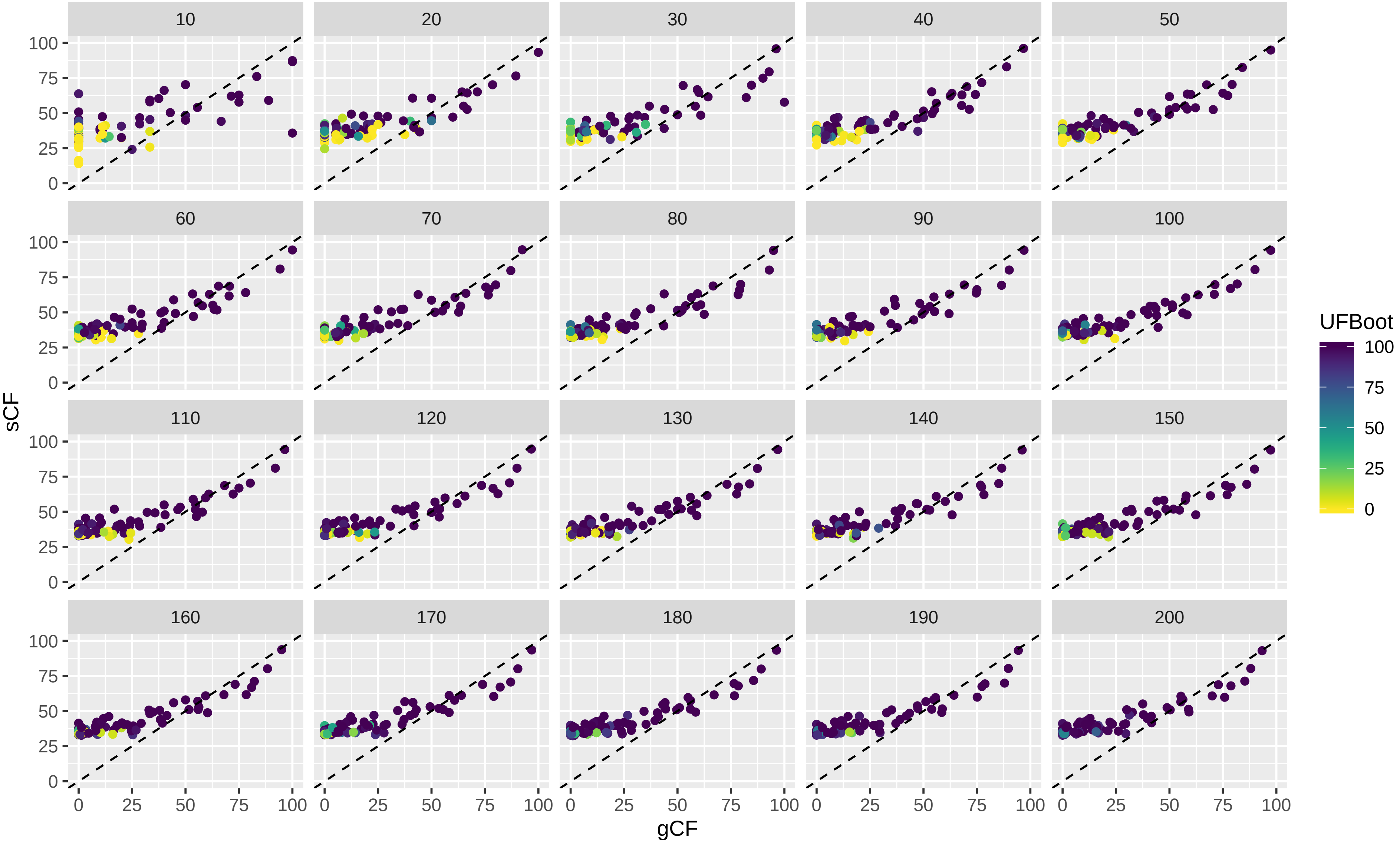

Supplementary Figure 3B

Dataset: Cannon_2016

The value of each variable (where the variable name is shown in the panel header) by number of loci used in the analysis.
Each coloured line represents a single branch in the tree estimated from the 200 locus dataset.

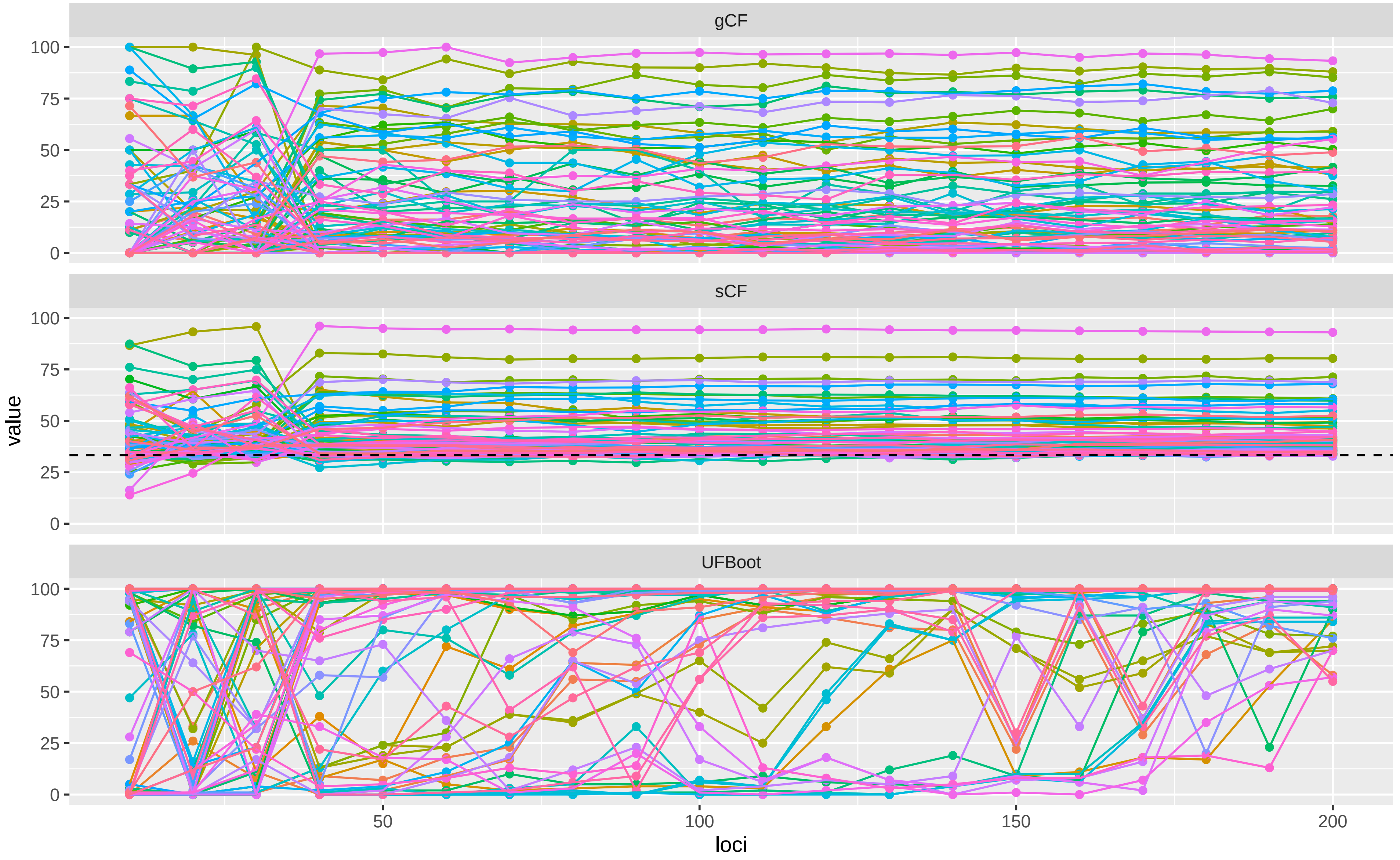

Supplementary Figure 3C

Dataset: Cannon_2016

The proportion of CF or Bootstrap values that are 100% (y axis) versus the number of loci used in the analysis (x axis).

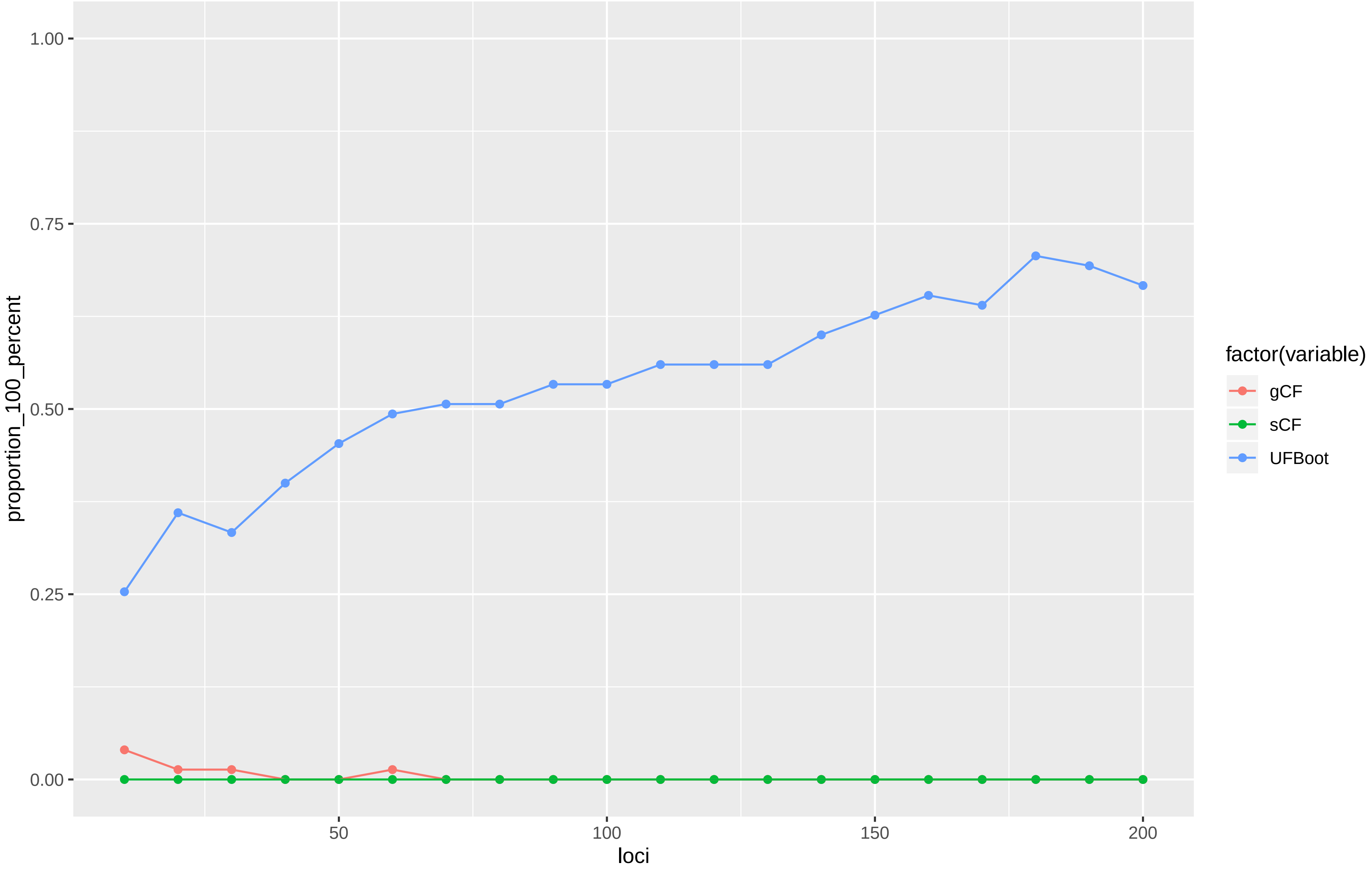

Supplementary Figure 4A

Dataset: Jarvis_2015

A scatterplot of gCF vs sCF values, coloured by UFBoot values. Each point represents a single branch in the tree calculated from the 200 locus dataset. Each panel represents an analysis in which the concordance factors and bootstraps were calculated from the number of loci specified in the panel header. The dashed line is the 1:1 line

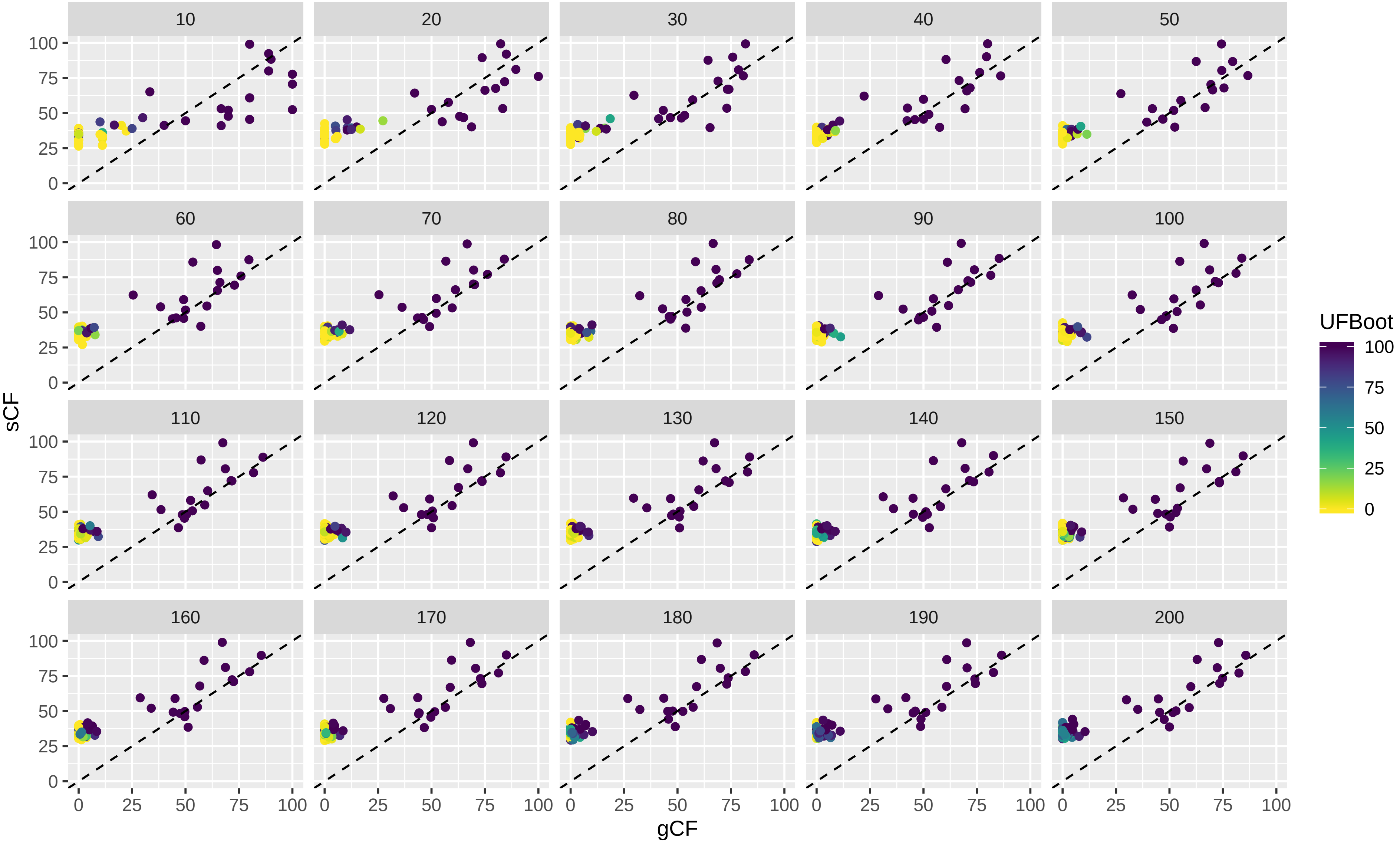

Supplementary Figure 4B

Dataset: Jarvis_2015
The value of each variable (where the variable name is shown in the panel header) by number of loci used in the analysis.
Each coloured line represents a single branch in the tree estimated from the 200 locus dataset.

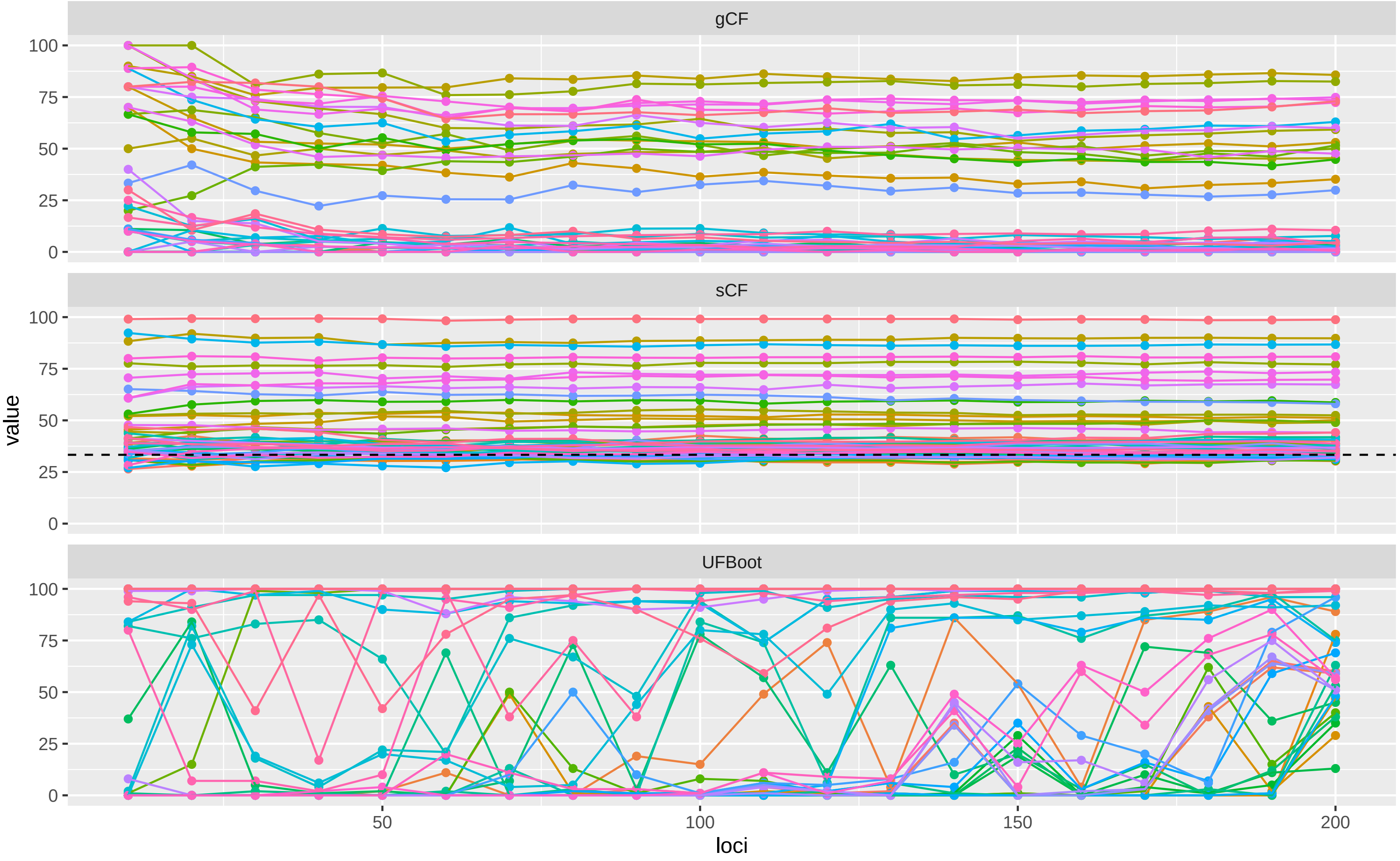

#### Supplementary Figure 4C

Dataset: Jarvis_2015

The proportion of CF or Bootstrap values that are 100% (y axis) versus the number of loci used in the analysis (x axis).

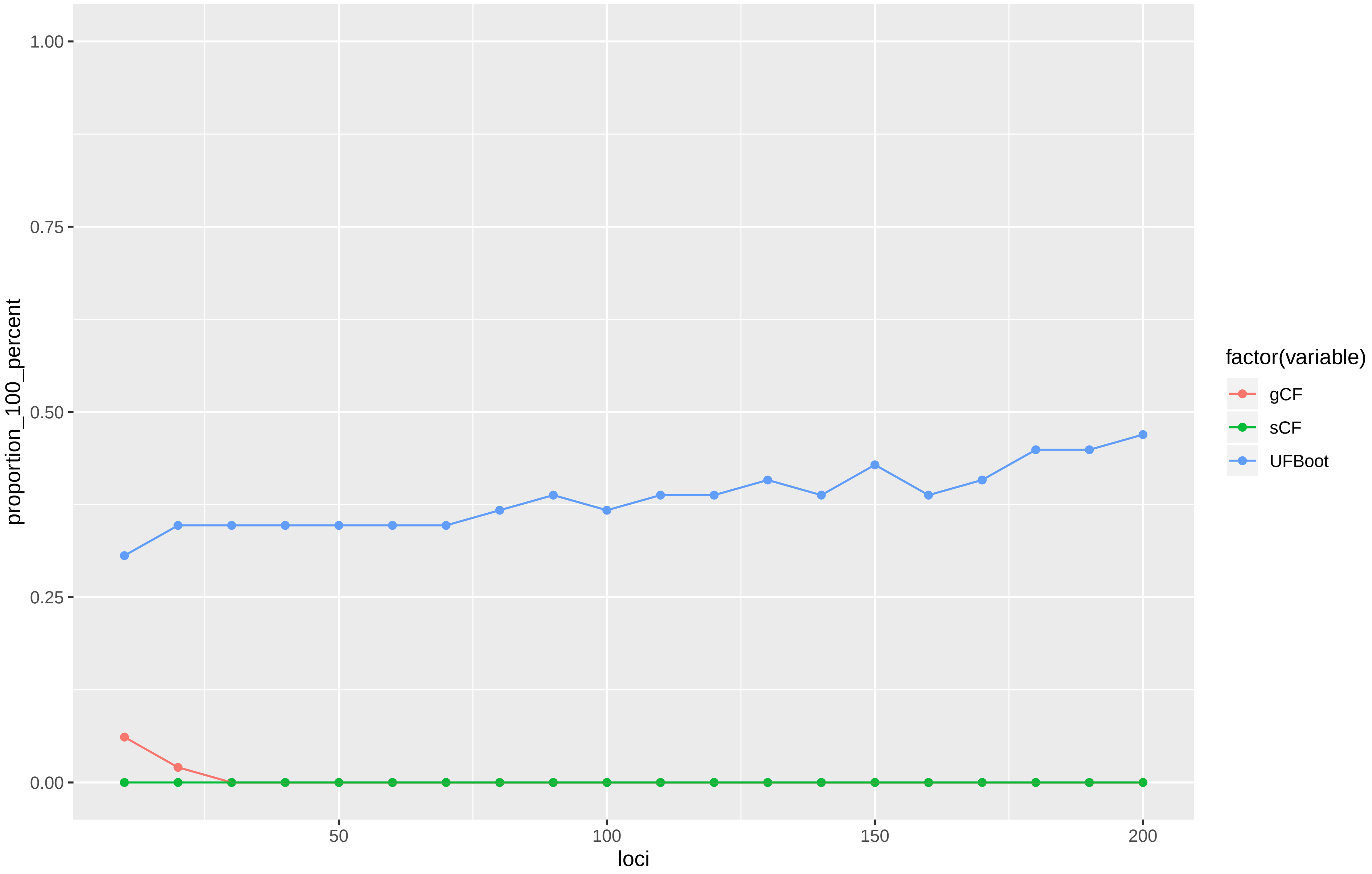

Supplementary Figure 5A

Dataset: Misof_2014
A scatterplot of gCF vs sCF values, coloured by UFBoot values. Each point represents a single branch in the tree calculated from the 200 locus dataset. Each panel represents an analysis in which the concordance factors and bootstraps were calculated from the number of loci specified in the panel header. The dashed line is the 1:1 line

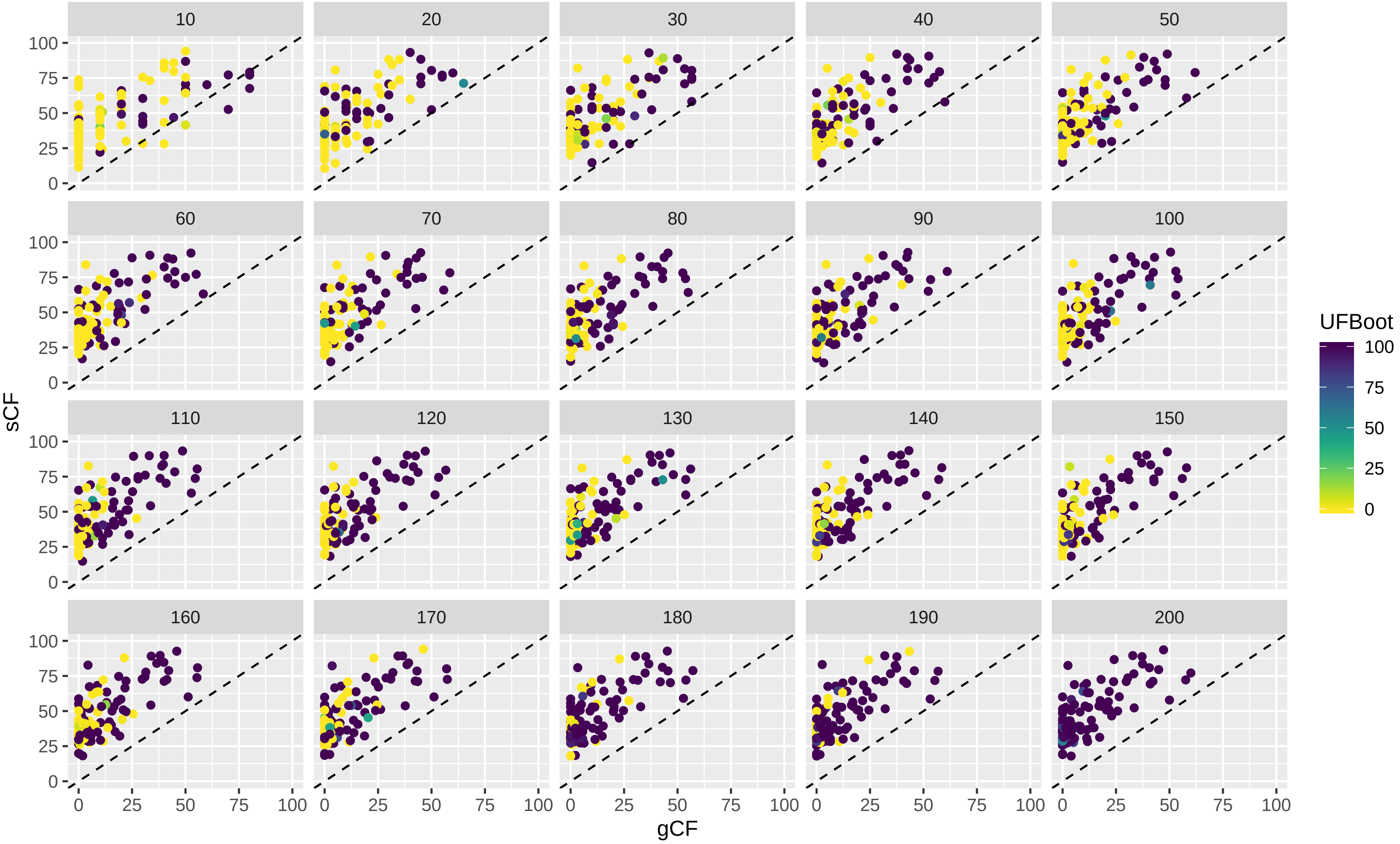

Supplementary Figure 5B

Dataset: Misof_2014
The value of each variable (where the variable name is shown in the panel header) by number of loci used in the analysis.
Each coloured line represents a single branch in the tree estimated from the 200 locus dataset.

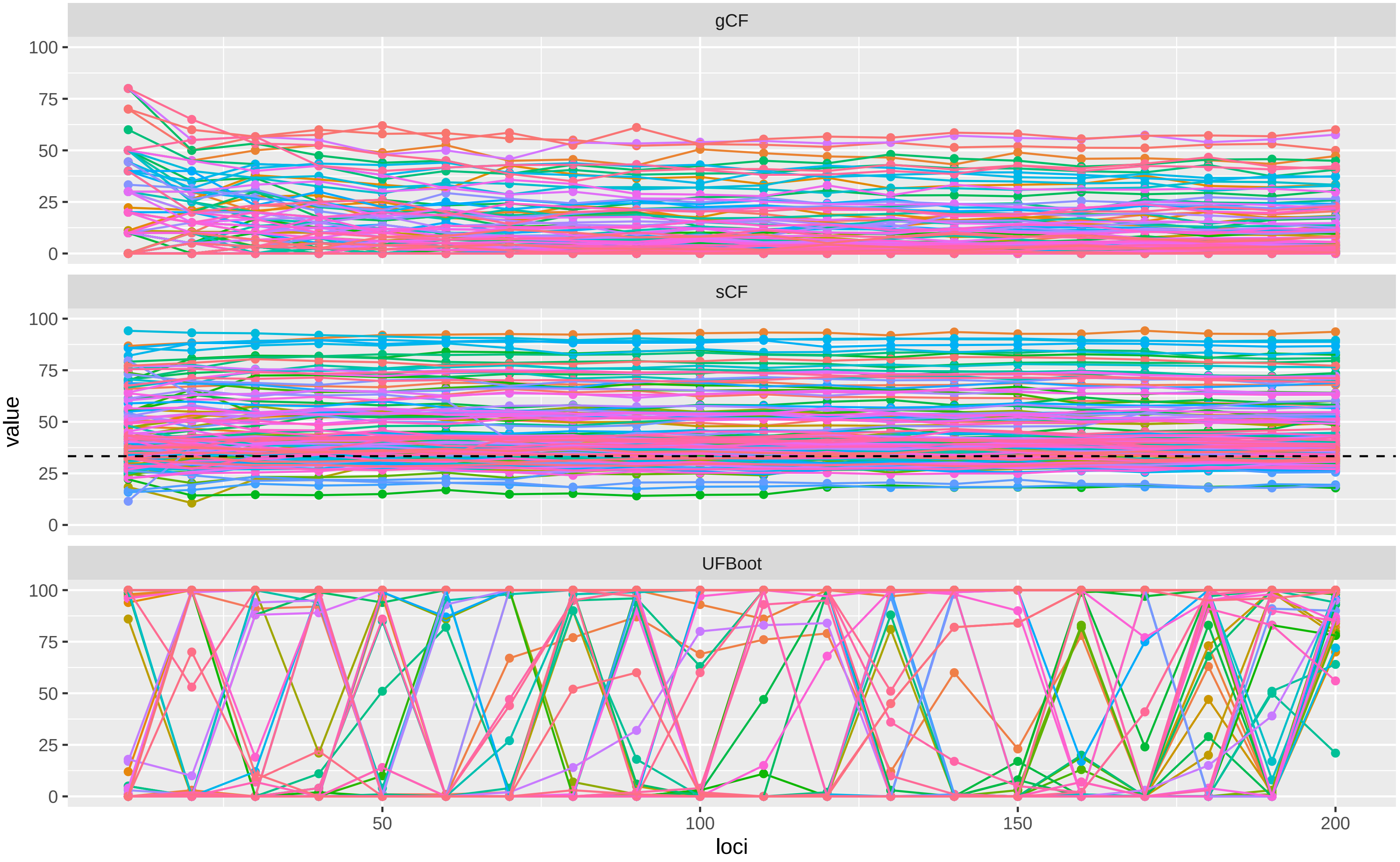

Supplementary Figure 5C

Dataset: Misof_2014

The proportion of CF or Bootstrap values that are 100% (y axis) versus the number of loci used in the analysis (x axis).

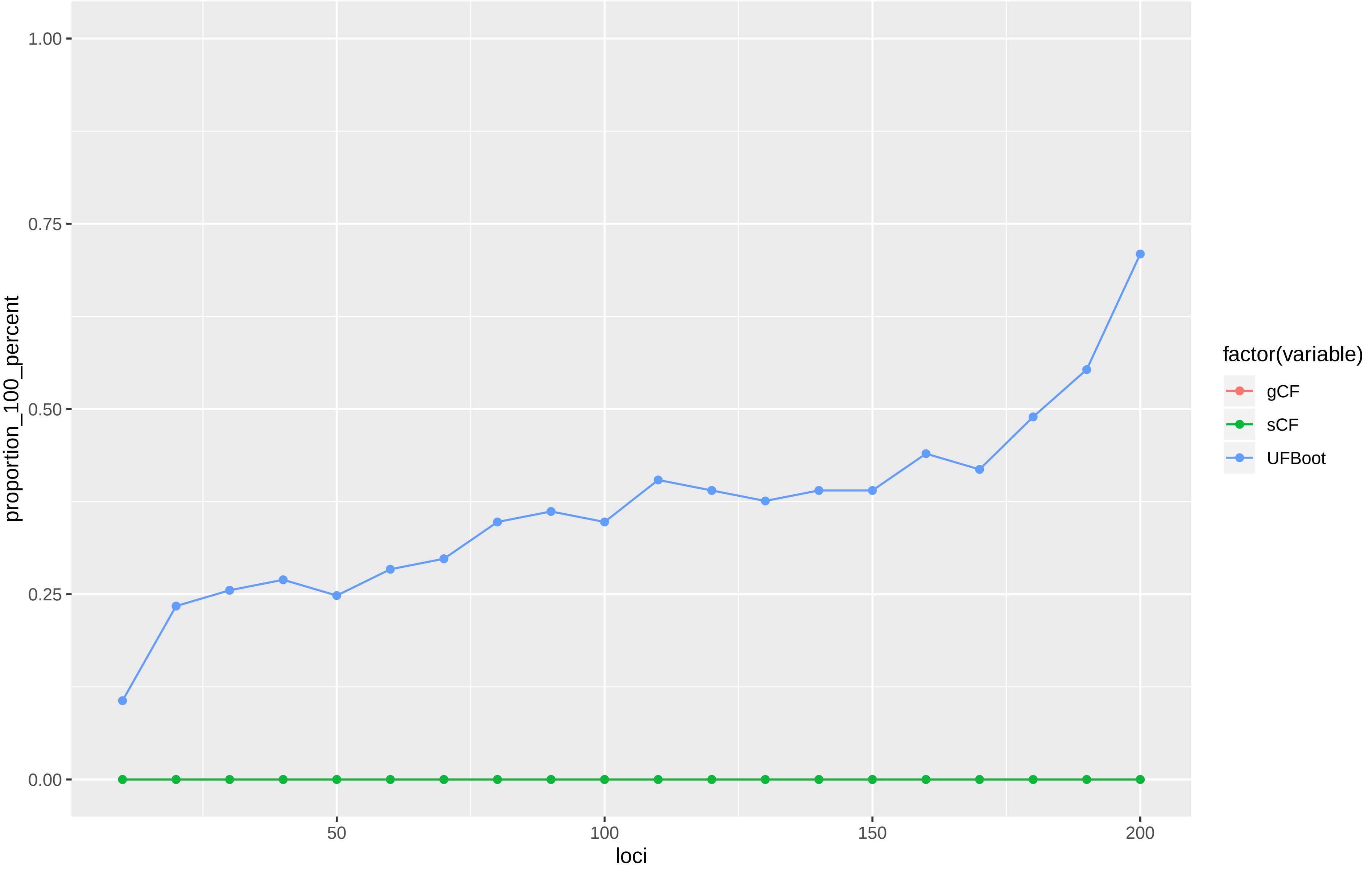

### Supplementary Figure 6A

Dataset: Ran_2018_aa

A scatterplot of gCF vs sCF values, coloured by UFBoot values. Each point represents a single branch in the tree calculated from the 200 locus dataset. Each panel represents an analysis in which the concordance factors and bootstraps were calculated from the number of loci specified in the panel header. The dashed line is the 1:1 line

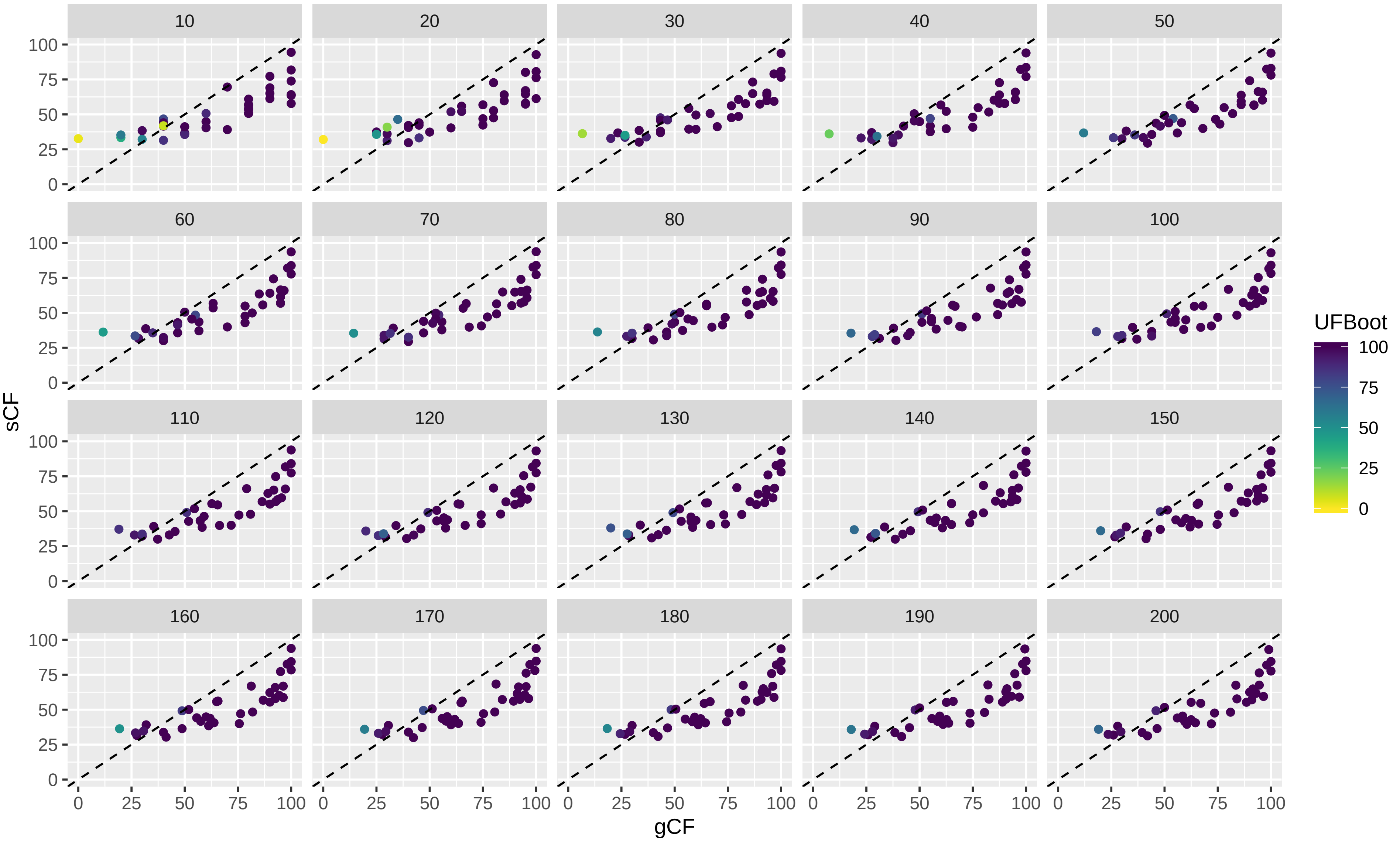

### Supplementary Figure 6B

Dataset: Ran_2018_aa
The value of each variable (where the variable name is shown in the panel header) by number of loci used in the analysis.
Each coloured line represents a single branch in the tree estimated from the 200 locus dataset.

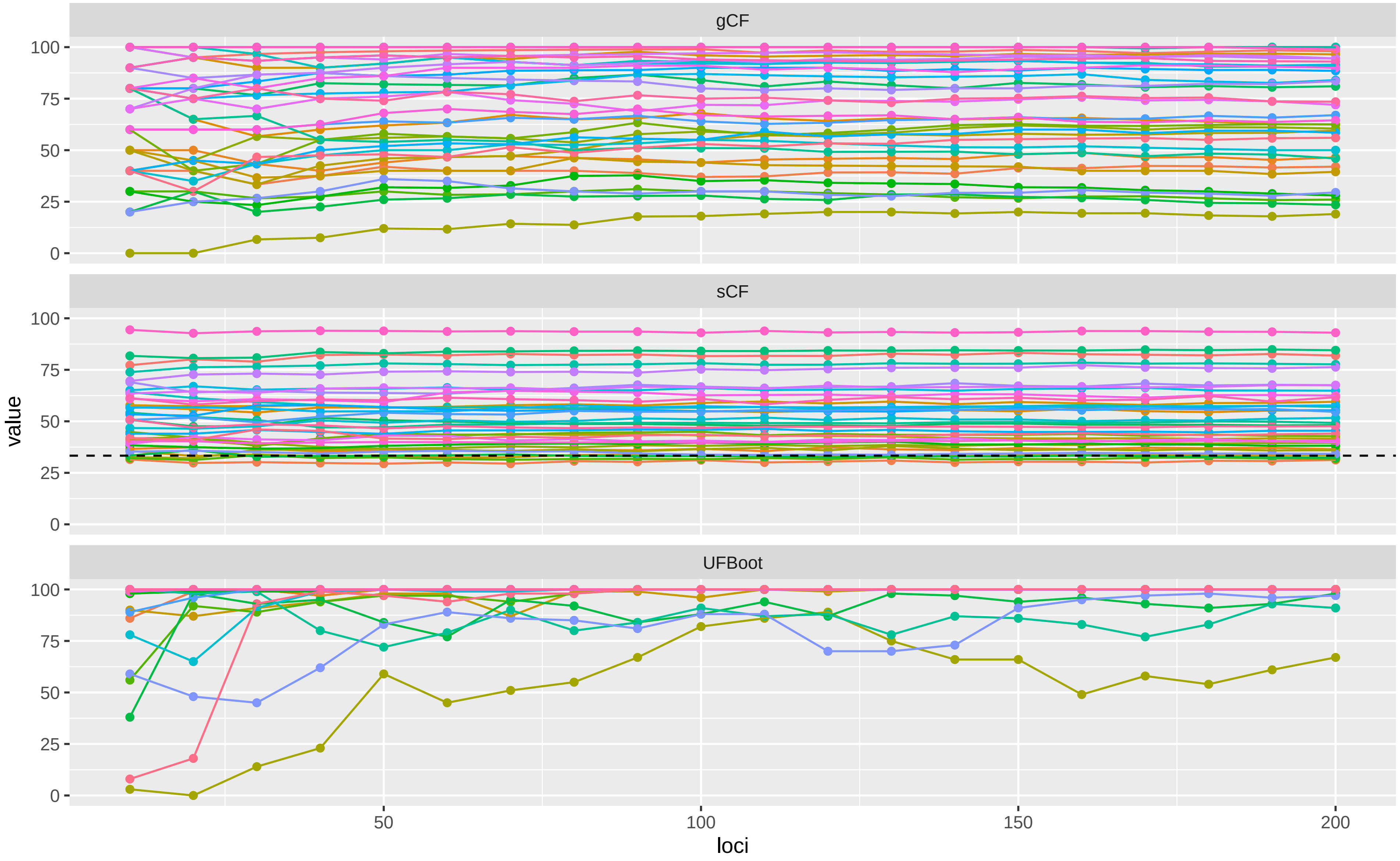

Supplementary Figure 6C

Dataset: Ran_2018_aa

The proportion of CF or Bootstrap values that are 100% (y axis) versus the number of loci used in the analysis (x axis).

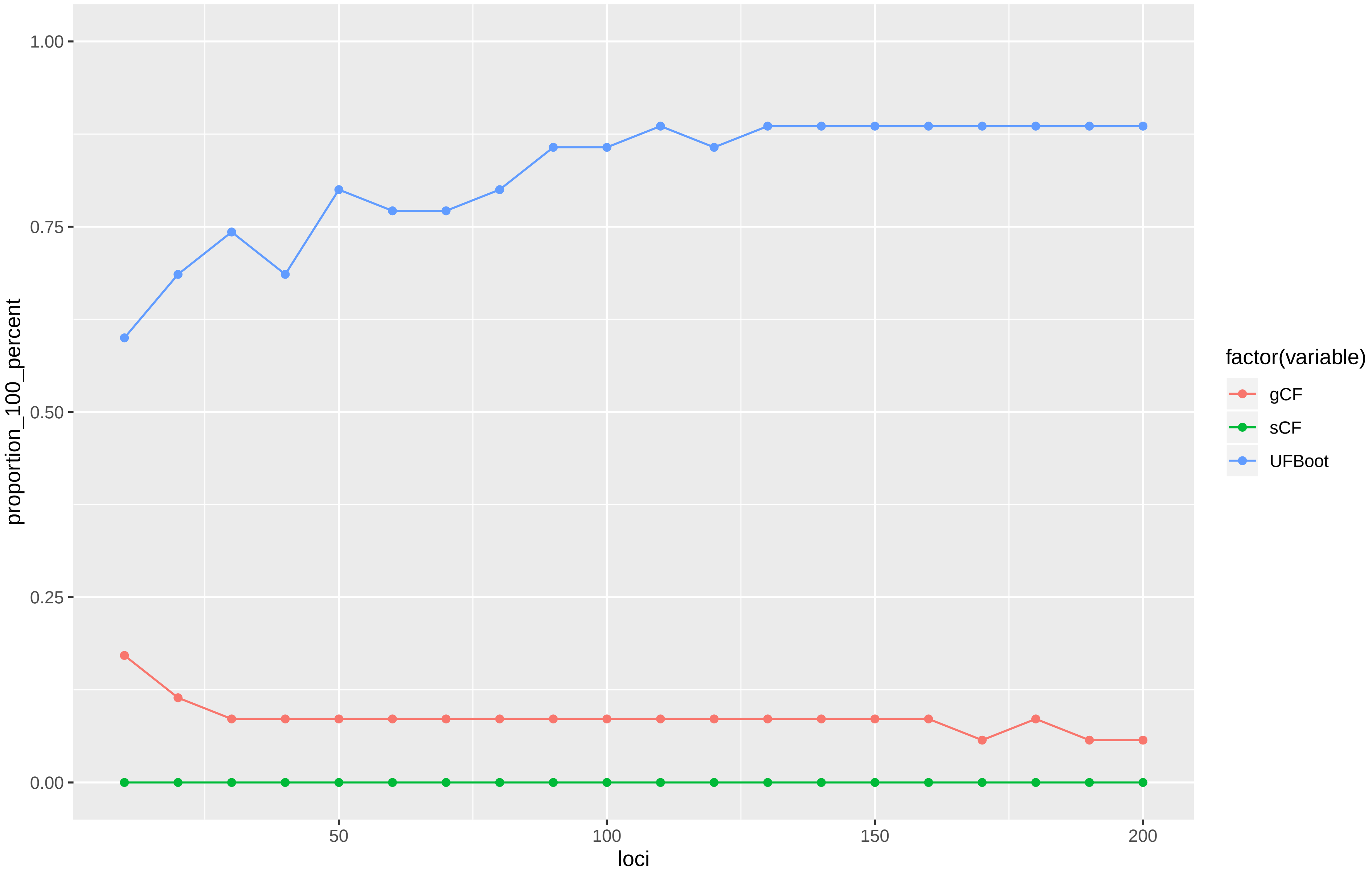

Supplementary Figure 7A

Dataset: Ran_2018_dna
A scatterplot of gCF vs sCF values, coloured by UFBoot values. Each point represents a single branch in the tree calculated from the 200 locus dataset. Each panel represents an analysis in which the concordance factors and bootstraps were calculated from the number of loci specified in the panel header. The dashed line is the 1:1 line

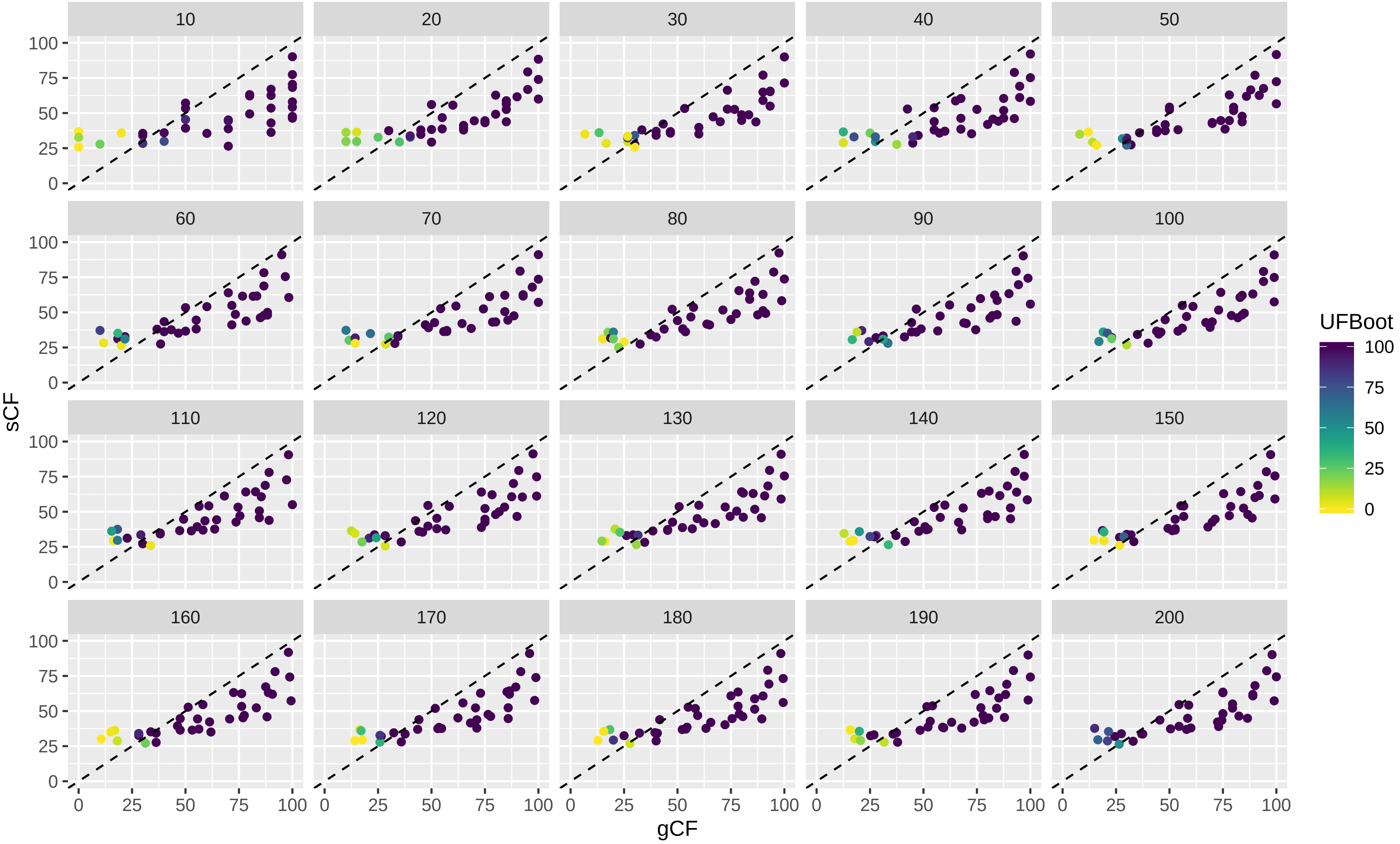

### Supplementary Figure 7B

Dataset: Ran_2018_dna
The value of each variable (where the variable name is shown in the panel header) by number of loci used in the analysis.
Each coloured line represents a single branch in the tree estimated from the 200 locus dataset.

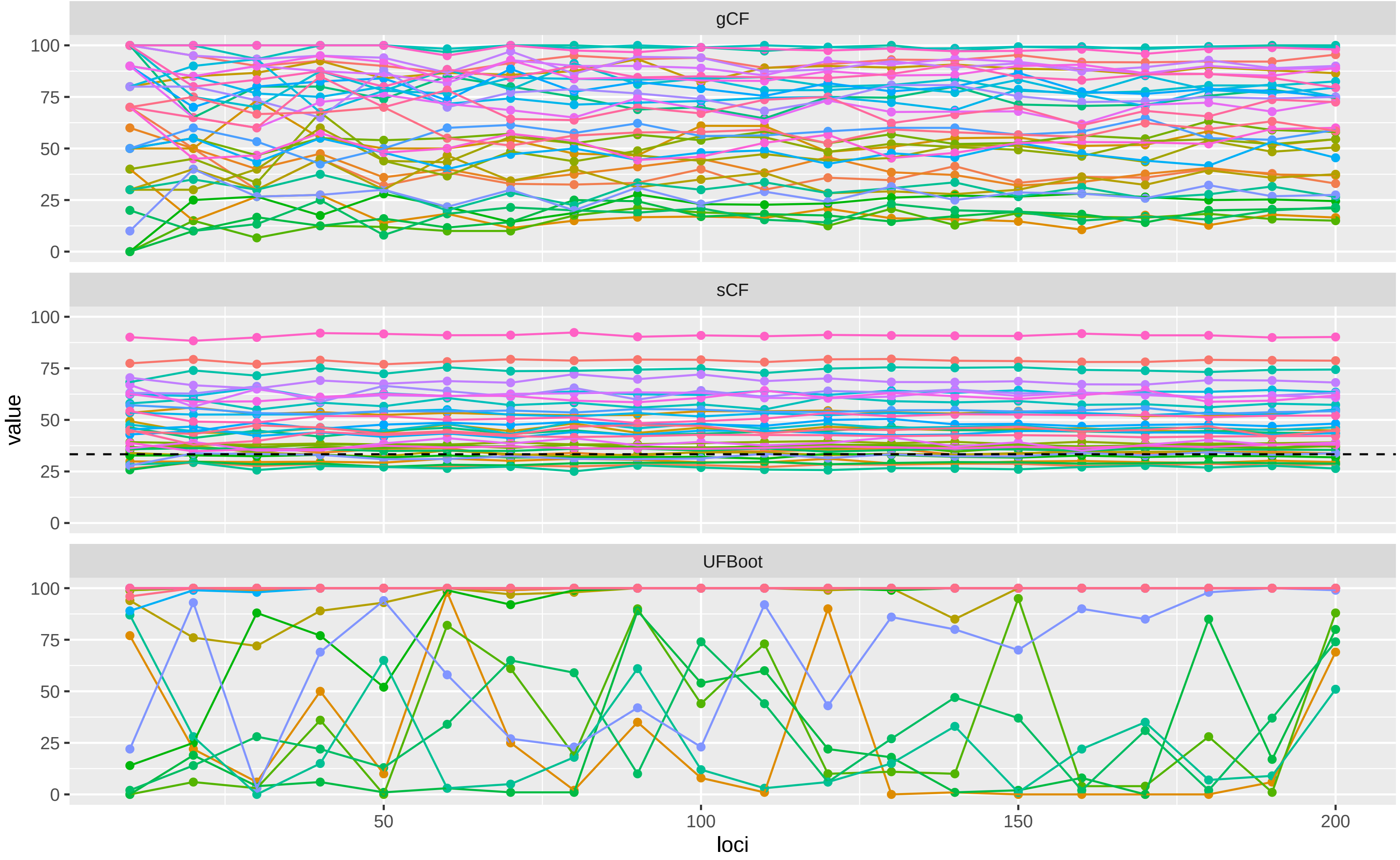

Supplementary Figure 7C

Dataset: Ran_2018_dna
The proportion of CF or Bootstrap values that are 100% (y axis) versus the number of loci used in the analysis (x axis).

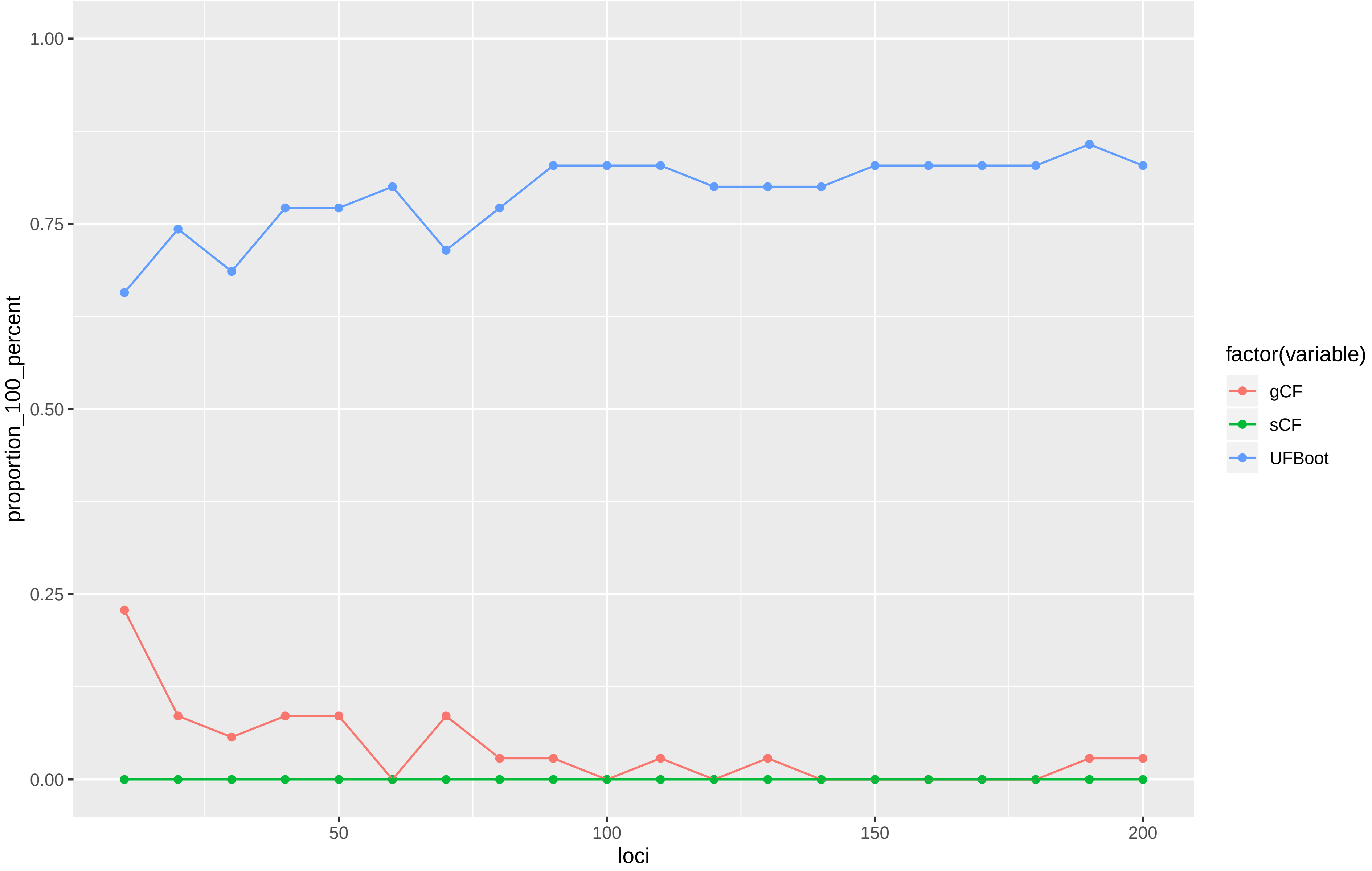

Supplementary Figure 8A

Dataset: Rodriguez_2018
A scatterplot of gCF vs sCF values, coloured by UFBoot values. Each point represents a single branch in the tree calculated from the 200 locus dataset. Each panel represents an analysis in which the concordance factors and bootstraps were calculated from the number of loci specified in the panel header. The dashed line is the 1:1 line

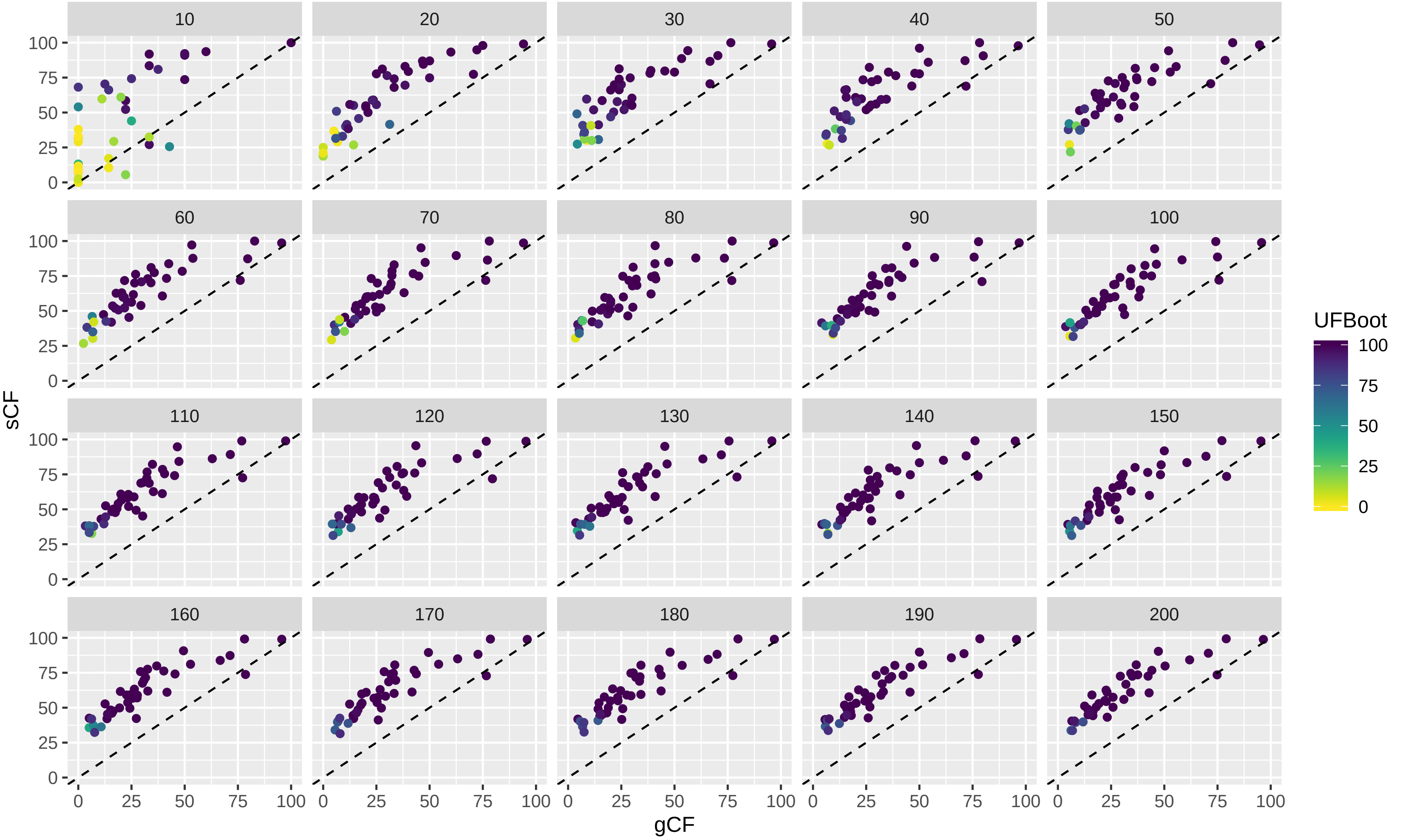

### Supplementary Figure 8B

Dataset: Rodriguez_2018
The value of each variable (where the variable name is shown in the panel header) by number of loci used in the analysis.
Each coloured line represents a single branch in the tree estimated from the 200 locus dataset.

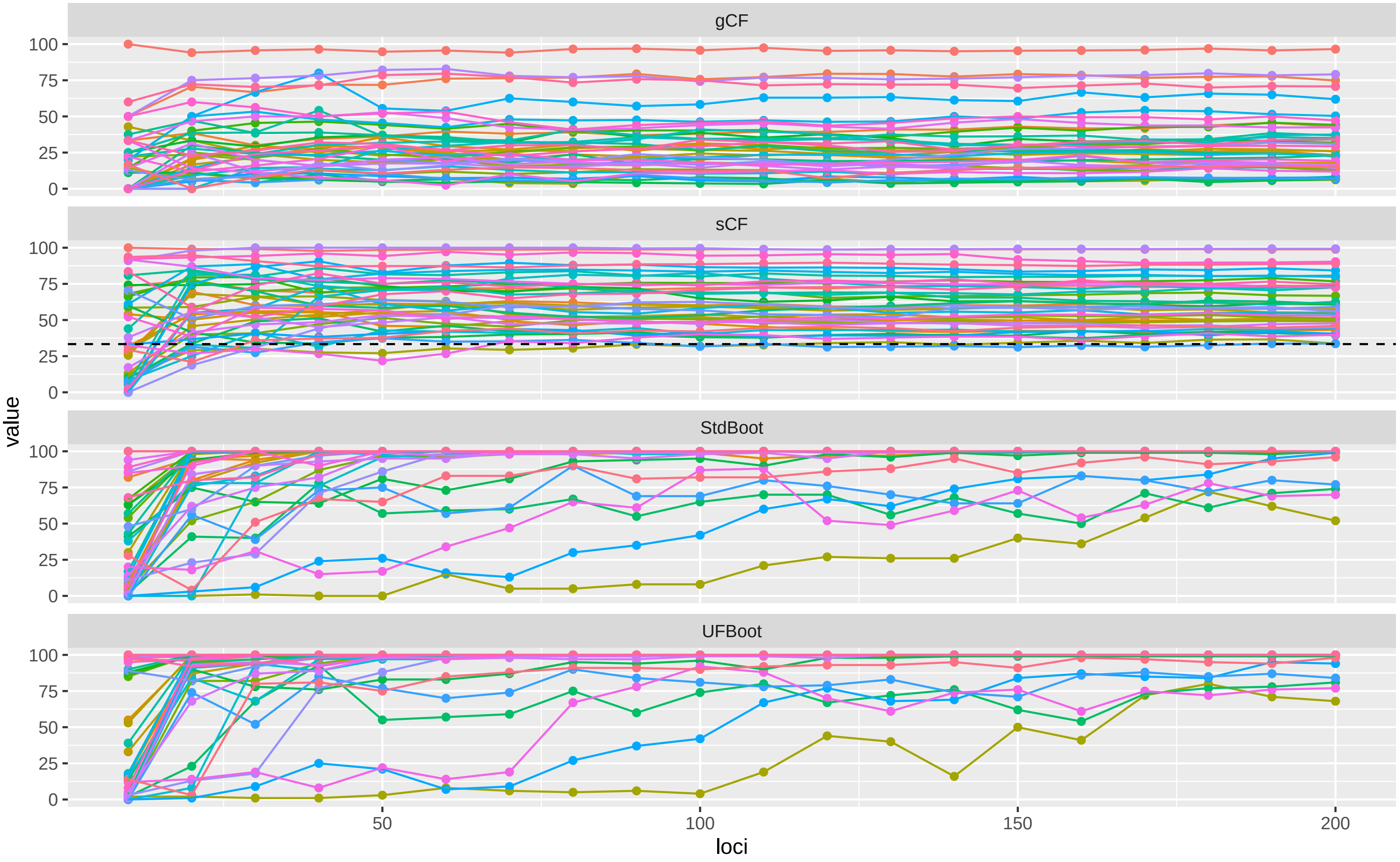

Supplementary Figure 8C

Dataset: Rodriguez_2018
The proportion of CF or Bootstrap values that are 100% (y axis) versus the number of loci used in the analysis (x axis).

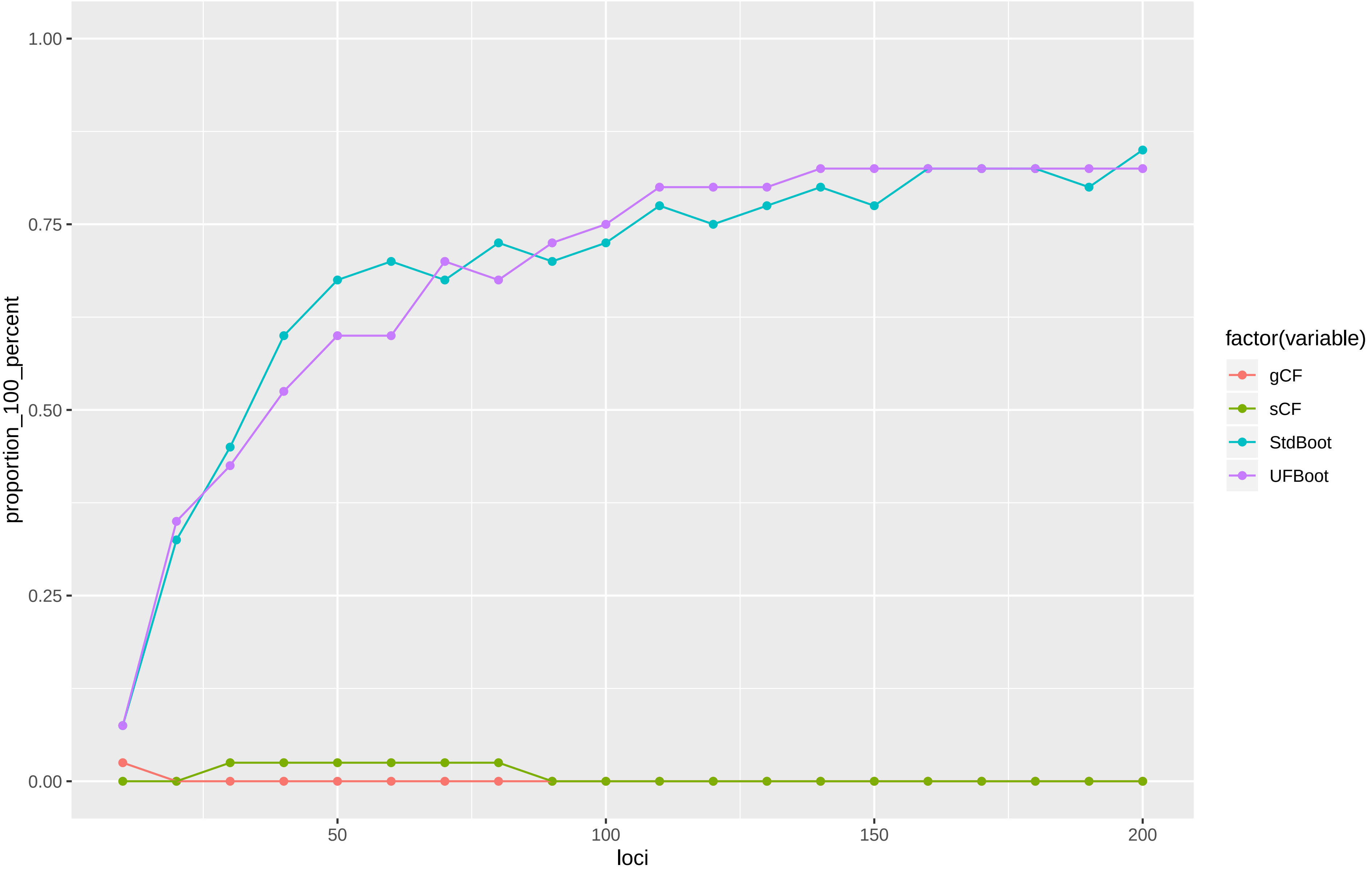

Supplementary Figure 8D

Dataset: Rodriguez_2018
A scatterplot of UFBoot vs. standard (i.e. Felsenstein) bootstrap values.
Each panel represents a different analysis using a number of loci specified in the panel header.
The dashed line is the 1:1 line

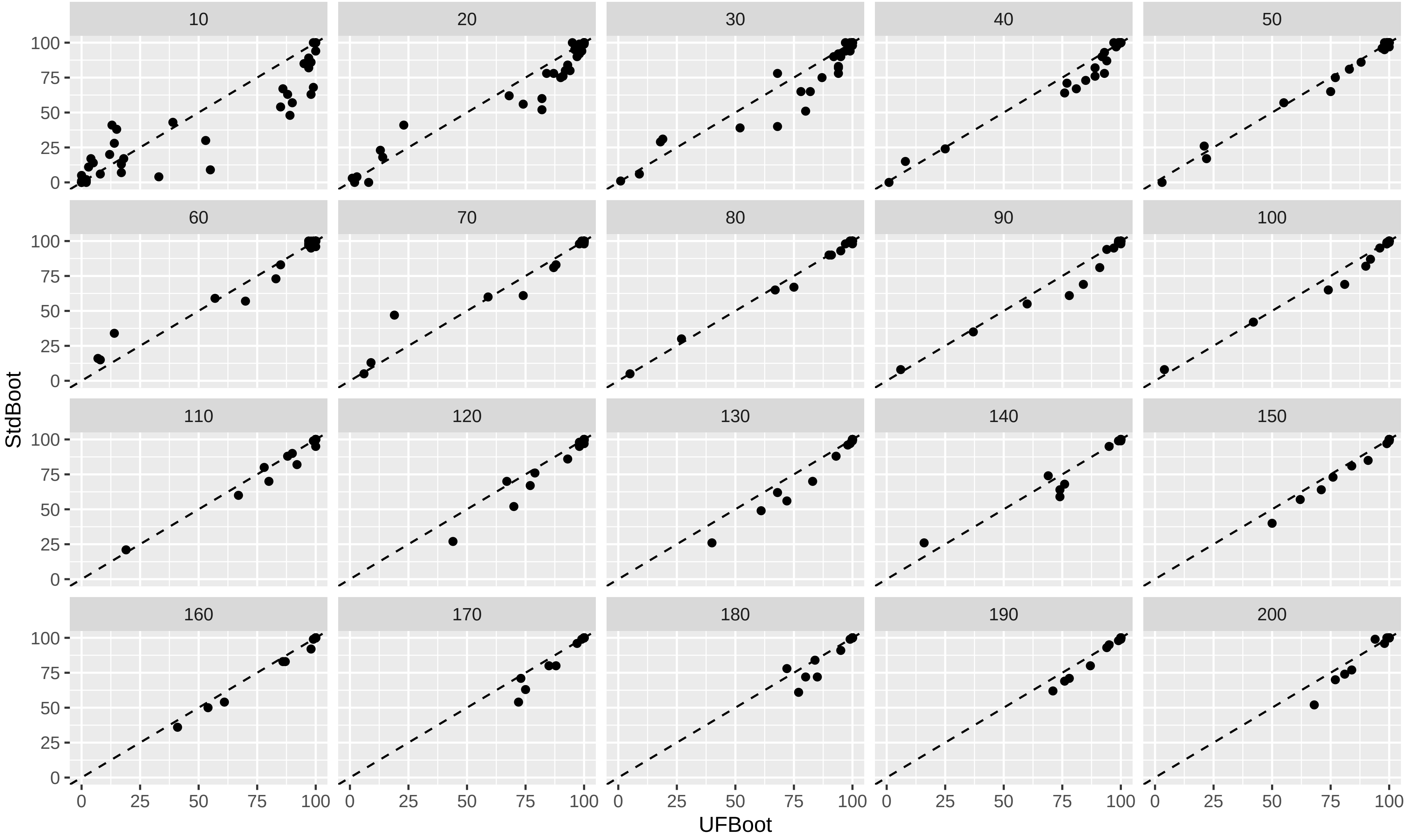

### Supplementary Figure 9A

Dataset: Wu_2018_aa
A scatterplot of gCF vs sCF values, coloured by UFBoot values. Each point represents a single branch in the tree calculated from the 200 locus dataset. Each panel represents an analysis in which the concordance factors and bootstraps were calculated from the number of loci specified in the panel header. The dashed line is the 1:1 line

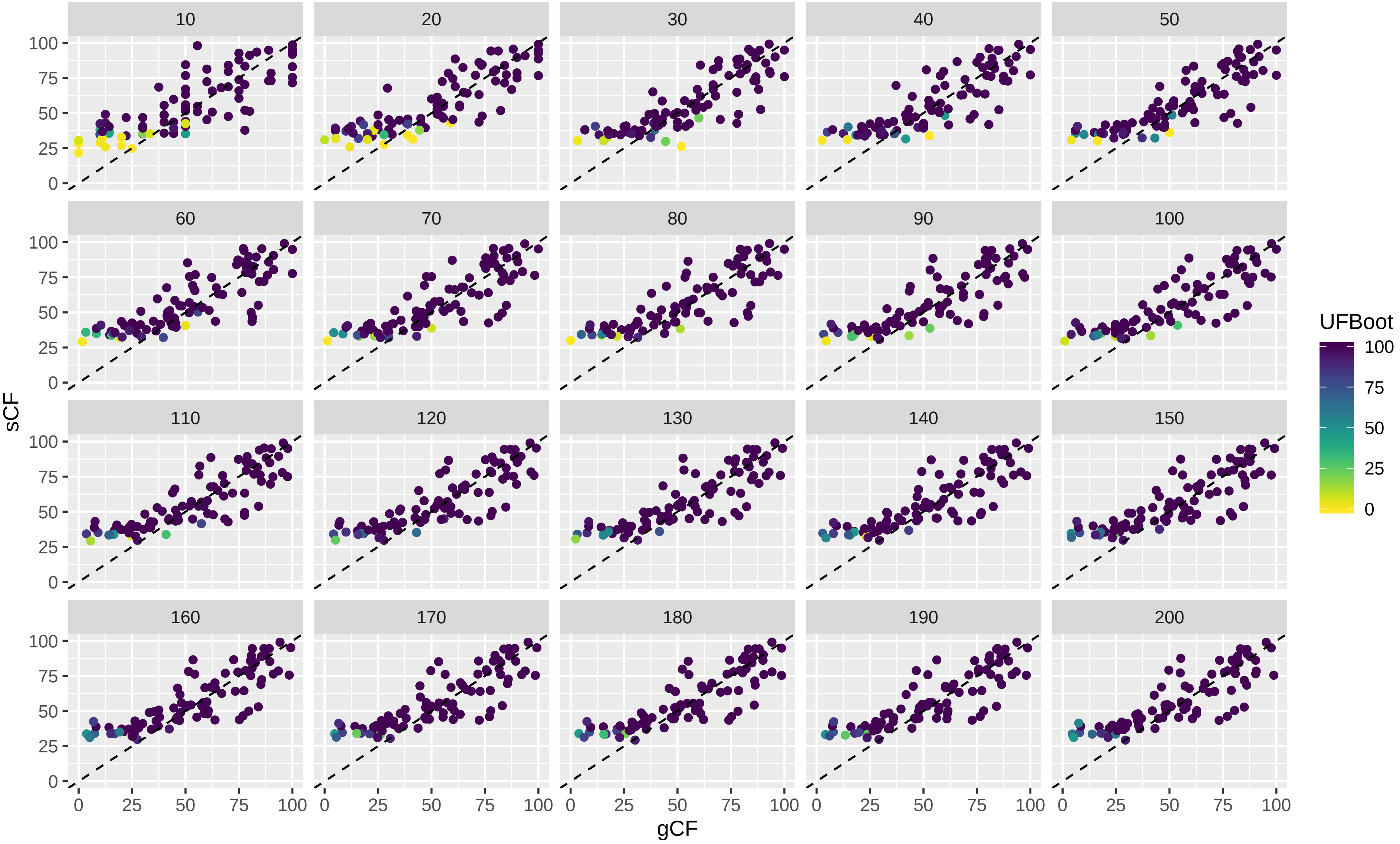

Supplementary Figure 9B

Dataset: Wu_2018_aa

The value of each variable (where the variable name is shown in the panel header) by number of loci used in the analysis.
Each coloured line represents a single branch in the tree estimated from the 200 locus dataset.

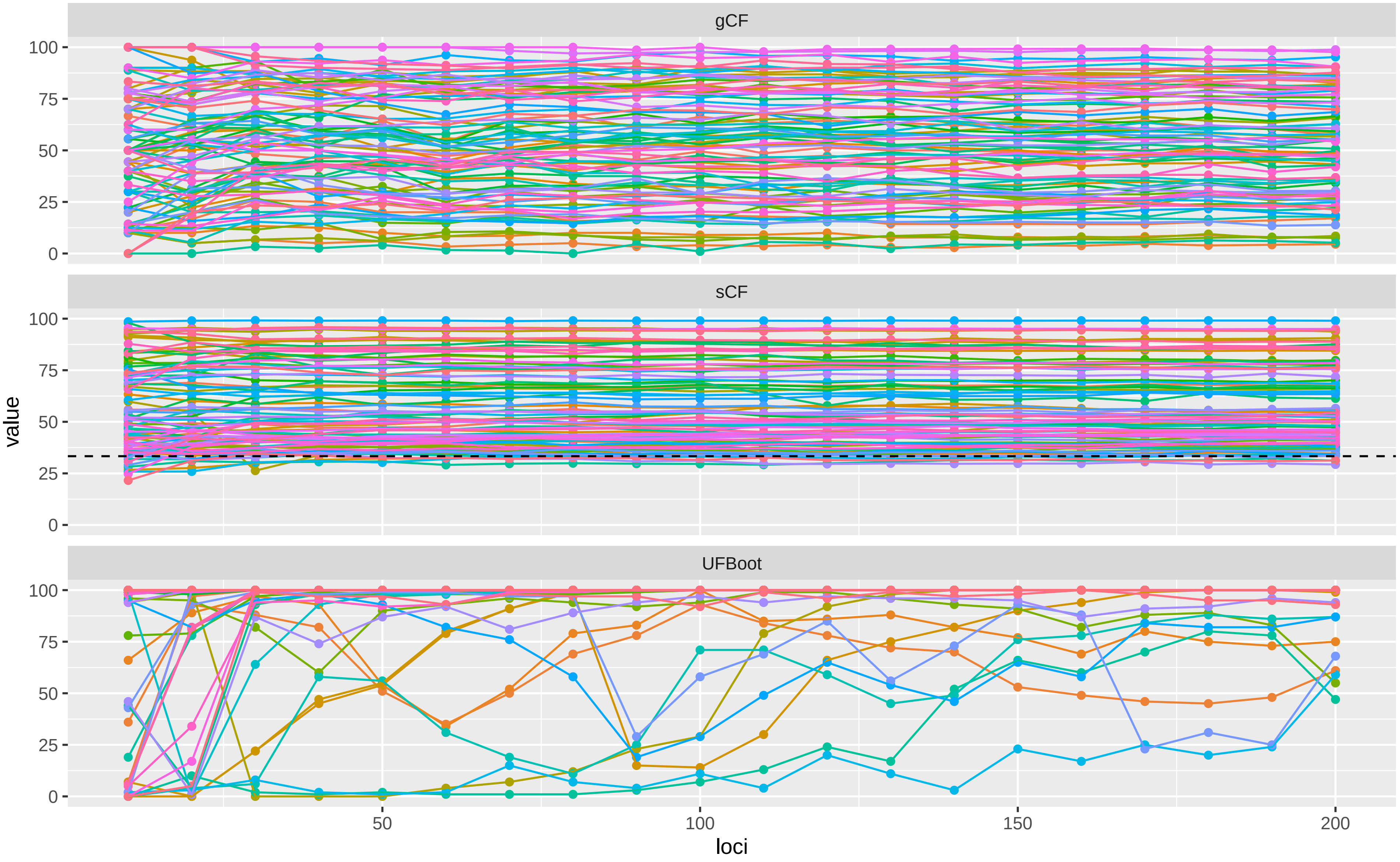

Supplementary Figure 9C

Dataset: Wu_2018_aa
The proportion of CF or Bootstrap values that are 100% (y axis) versus the number of loci used in the analysis (x axis).

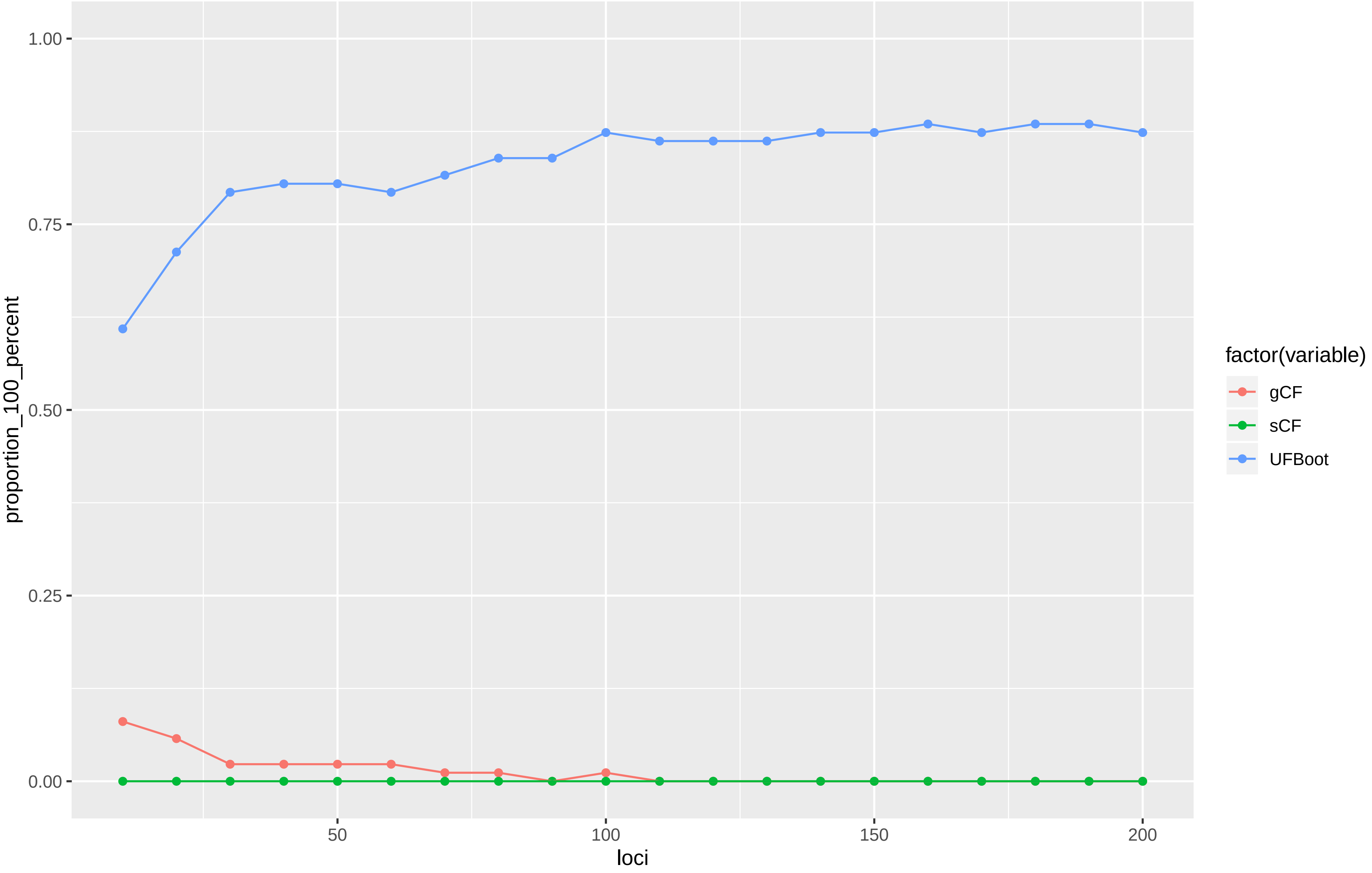

#### Supplementary Figure 10A

Dataset: Wu_2018_dna

A scatterplot of gCF vs sCF values, coloured by UFBoot values. Each point represents a single branch in the tree calculated from the 200 locus dataset. Each panel represents an analysis in which the concordance factors and bootstraps were calculated from the number of loci specified in the panel header. The dashed line is the 1:1 line

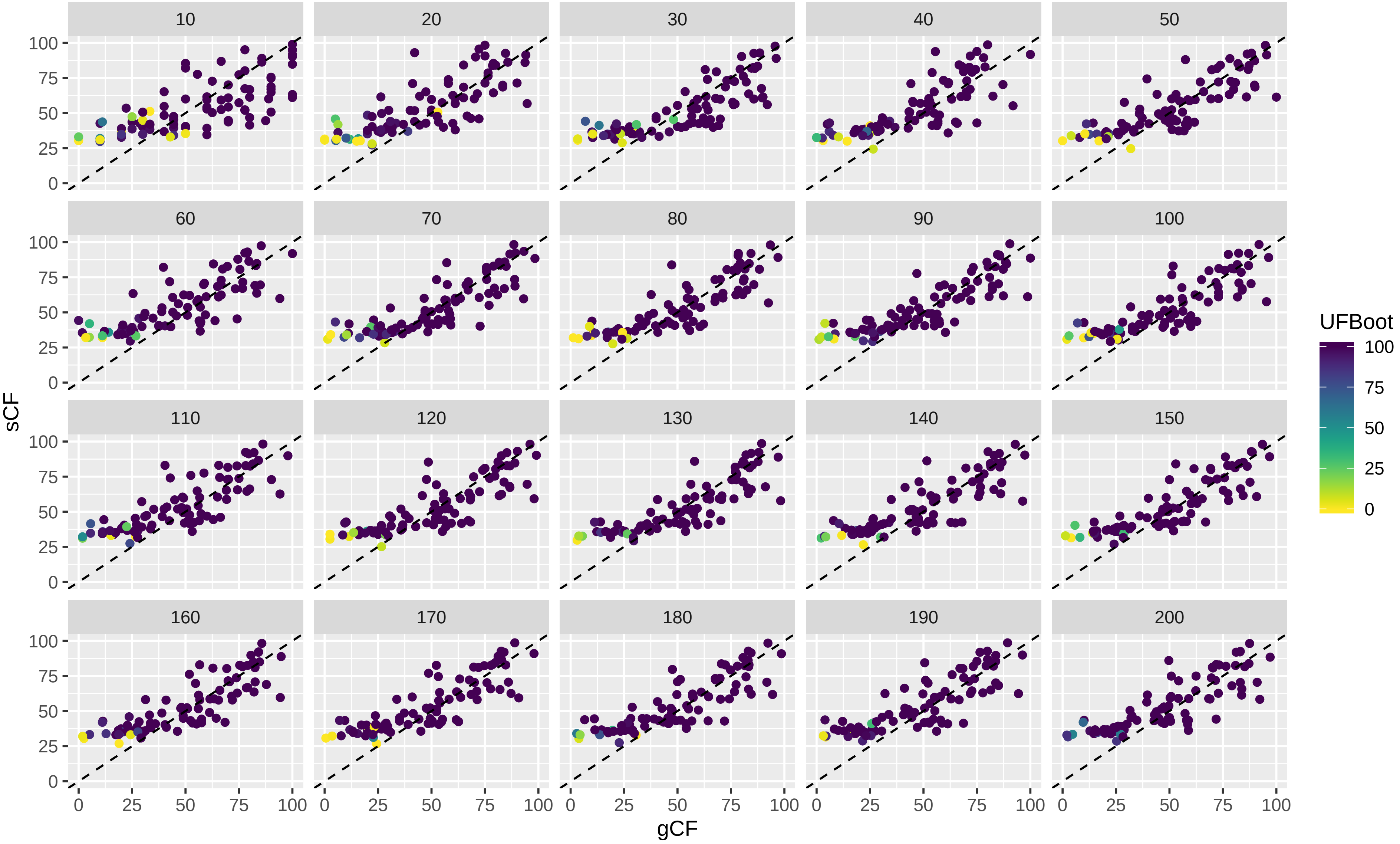

Supplementary Figure 10B

Dataset: Wu_2018_dna
The value of each variable (where the variable name is shown in the panel header) by number of loci used in the analysis.
Each coloured line represents a single branch in the tree estimated from the 200 locus dataset.

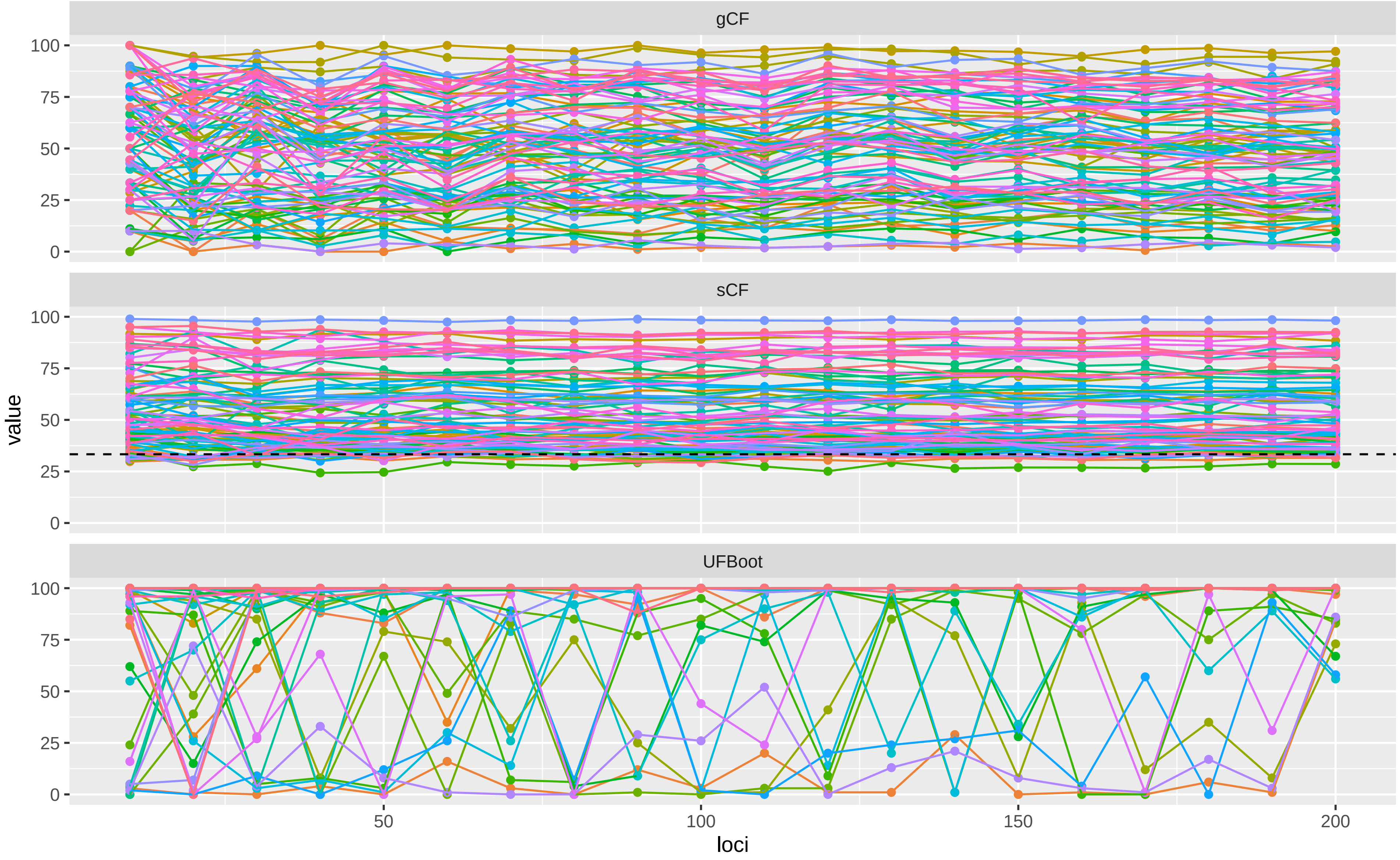

### Supplementary Figure 10C

Dataset: Wu_2018_dna

The proportion of CF or Bootstrap values that are 100% (y axis) versus the number of loci used in the analysis (x axis).

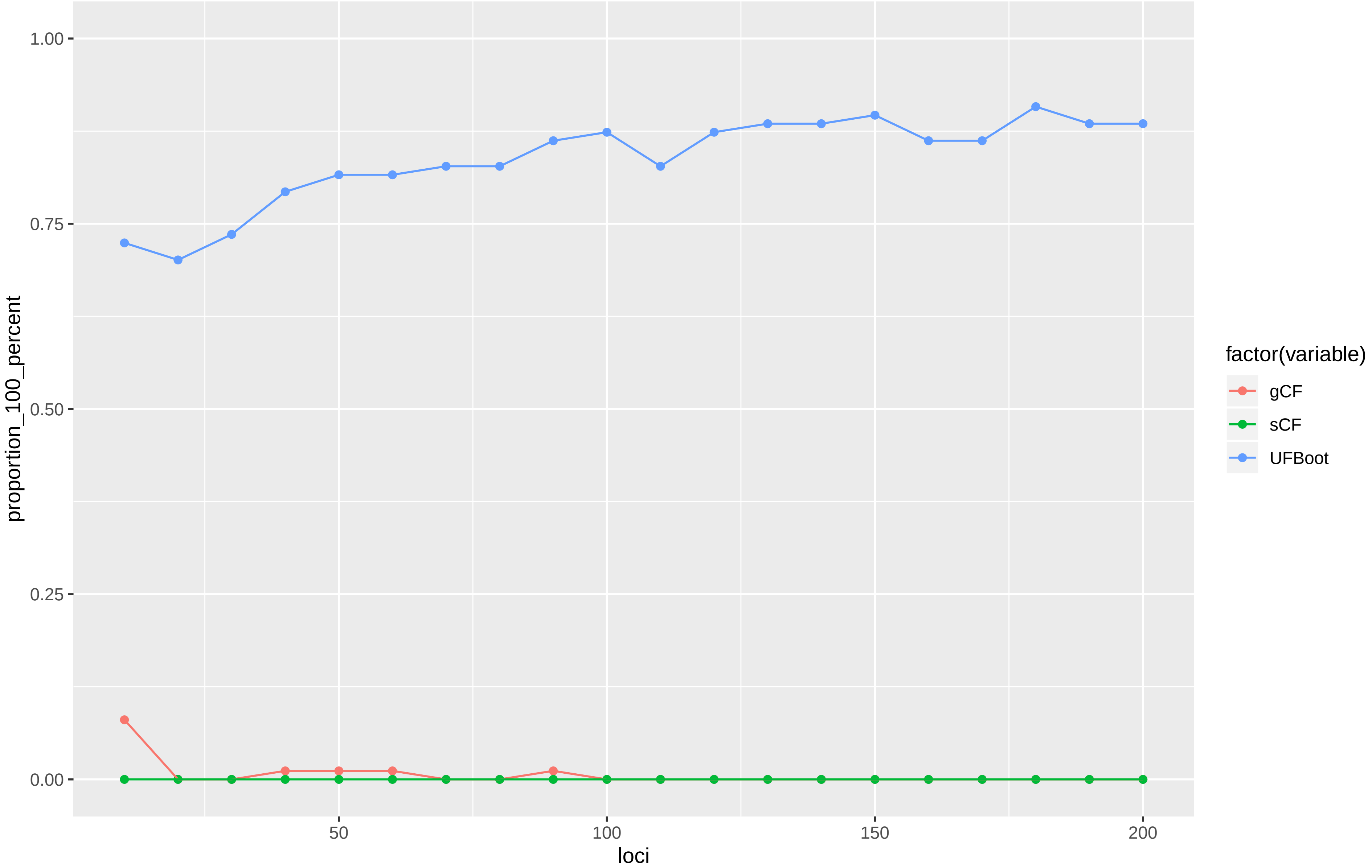
